# Early immunomodulatory program triggered by pro-tolerogenic *Bifidobacterium pseudolongum* drives cardiac transplant outcomes

**DOI:** 10.1101/2022.10.13.511915

**Authors:** Samuel J Gavzy, Allison Kensiski, Vikas Saxena, Ram Lakhan, Lauren Hittle, Long Wu, Jegan Iyyathurai, Hima Dhakal, Zachariah L Lee, Lushen Li, Young S Lee, Tianshu Zhang, Hnin W Lwin, Marina W Shirkey, Christina M Paluskievicz, Wenji Piao, Emmanuel F Mongodin, Bing Ma, Jonathan S Bromberg

## Abstract

**Background:** Despite ongoing improvements in regimens to prevent allograft rejection, most cardiac and other organ grafts eventually succumb to chronic vasculopathy, interstitial fibrosis, or endothelial changes, and eventually graft failure. The events leading to chronic rejection are still poorly understood and the gut microbiota is a known driving force in immune dysfunction. We previously showed that gut microbiota dysbiosis profoundly influences the outcome of vascularized cardiac allografts and subsequently identified biomarker species associated with these differential graft outcomes.

**Methods:** In this study, we further detailed the multifaceted immunomodulatory properties of pro-tolerogenic and pro-inflammatory bacterial species over time, using our clinically relevant model of allogenic heart transplantation.

**Results:** In addition to tracing longitudinal changes in the recipient gut microbiome over time, we observed that *Bifidobacterium pseudolongum (Bifido)* induced an early anti-inflammatory phenotype within 7 days, while *Desulfovibrio desulfuricans (Desulfo)* resulted in a pro-inflammatory phenotype, defined by alterations in leukocyte distribution and lymph node (LN) structure. Indeed, *in vitro* results showed that *Bifido* and *Desulfo* acted directly on primary innate immune cells. However, by 40 days after treatment, these two bacterial strains were associated with mixed effects in their impact on LN architecture and immune cell composition and loss of colonization within gut microbiota, despite protection of allografts from inflammation with *Bifido* treatment.

**Conclusions:** These dynamic effects suggest a critical role for early microbiota-triggered immunological events such as innate immune cell engagement, T cell differentiation, and LN architectural changes in the subsequent modulation of pro-tolerant versus pro-inflammatory immune responses in organ transplant recipients.

## INTRODUCTION

One-year survival rates in solid organ transplantation routinely exceed 90-95%; however, most cardiac and other types of allografts eventually succumb to chronic vascular, interstitial, or endothelial impairment beyond the first year, despite the many improvements over the past decades in organ preservation, recipient selection, immunosuppressive regimens and monitoring, and infection prophylaxis^1–7^. As a result, the long-term rate of decline of allograft function has changed little in the past 20 years^5,8^. The immunological events leading to chronic rejection are still poorly understood, with a resulting dearth of predictive diagnostic tools and interventions. Chronic rejection is only loosely associated with T and B cell immunity^2,3,9^, necessitating a novel approach to characterize recipient alloimmune responses.

Numerous studies have shown that the microbiota is a driving force in immune dysfunction and thus a potential diagnostic or therapeutic target^10–12^. The gut microbiota was shown to trigger pro-or anti-inflammatory signals that regulate local and systemic innate and adaptive immune homeostasis^13^. While changes in the microbiota following organ and bone marrow transplant have been well documented^14–25^, these studies have been largely associative. These observations provide a rationale for characterizing the microbiota and determining the precise mechanisms it employs to modulate immunity and affect chronic organ damage.

The immune system and microbiota have bidirectional interactions, and in transplantation, immunosuppression alters the immune system, while antibiotic (abx) and immunosuppression administration both alter the microbiota^26–29^. Disruption of the microbiota, for example through abx treatment, has been shown to impact graft survival. We previously demonstrated that both the gut microbiota as a whole or the responsible single bacteria (*Bifido* or *Desulfo* species) can profoundly influence the outcomes of vascularized cardiac allografts^30^. Oral gavage with a single anti-inflammatory strain of *Bifidobacterium pseudolongum* (*Bifido*), enriched in pregnant mouse (representing immune suppression) gut microbiome, resulted in significant improvement in long-term allograft survival, prevention of inflammation and fibrosis in the grafts, and balanced anti-inflammatory and homeostatic cytokine response from dendritic cells (DCs) and macrophages (MΦ)^30,31^. On the other hand, the single pro-inflammatory strain of *Desulfovibrio desulfuricans* (*Desulfo*), enriched in colitic mouse (representing immune inflammation) gut microbiome, triggered a greater inflammatory cytokine response *in vitro*^30^. *Bifido* also induced changes in lymph node (LN) structure, resulting in an increased ratio of laminin α4:α5 in the cortical ridge (CR), mechanistically associated with immunologic suppression and tolerance^30^. In comparison, *Desulfo* was associated with inflammation and induced diametrical effects with reduced laminin α4:α5 ratios^30^.

This study expands on our initial findings^30^ by further dissecting the molecular pathways involved in the interactions among specific immune cell populations, LNs, and selected members of the gut microbiota (i.e., *Bifido* and *Desulfo* species). Using our *in vivo* model combining single bacterial strain transfer (SBT), allogenic heart transplant, abx and immunosuppression treatment, we further detailed the dynamic immunomodulatory properties of pro-tolerogenic and pro-inflammatory bacteria on short and long term graft outcomes. We also characterized dynamic longitudinal alterations in recipient gut microbiomes following one time treatment with single bacterial species. These results demonstrate persistent changes in recipient alloimmunity despite eventual disappearance of transferred bacteria. The effect of *Bifido* on LN architecture as well immune cell composition and differentiation was most pronounced within 7 days of treatment, with the bacteria-driven effects becoming attenuated over time. Lastly, we demonstrate strain-specific effects of these bacteria on the activation profiles of primary myeloid cells *in vitro*. These dynamic changes highlight the importance of early innate immune cell engagement and LN architectural changes in the modulation of pro-tolerant versus pro-inflammatory immune responses in organ transplant recipients.

## RESULTS

### Early profound impact of bacterial gavage on microbiota with equilibration at later time points

In order to assess the stability of colonization by gavaged bacteria over time, mice were treated with antibiotics (abx) to deplete their normal microbiota, after which gavage was performed using wild type (WT) mouse whole stool samples, or single bacterial strain transfer (SBT) of murine-isolated *B. pseudolongum* UMB-MBP-001 (*Bifido* MD), or *D. desulfuricans* (*Desulfo*). Cardiac transplants were then performed and followed by daily low dose tacrolimus immunosuppression. The gut microbiota was characterized longitudinally using 16S rRNA gene amplicon sequencing over 40 days. The α diversity (within-sample diversity) of the microbial communities, as measured by Shannon diversity index^32^, showed that overall microbial diversity decreased within the first 2 weeks after *Bifido* and *Desulfo* SBT, with restoration of diversity at later time points **(Fig. 1a)**. The β-diversity (between-sample diversity) of the microbial communities, calculated using Jensen–Shannon divergence distance, demonstrated distinct separation of *Bifido* and *Desulfo* after SBT, while the gut microbiota composition profiles of all 3 treatment groups converged by week 2 **(Fig. 1b**). These results indicate the profound, yet transient impact of both *Bifido* and *Desulfo* on the gut microbiota.

**Figure 1.**
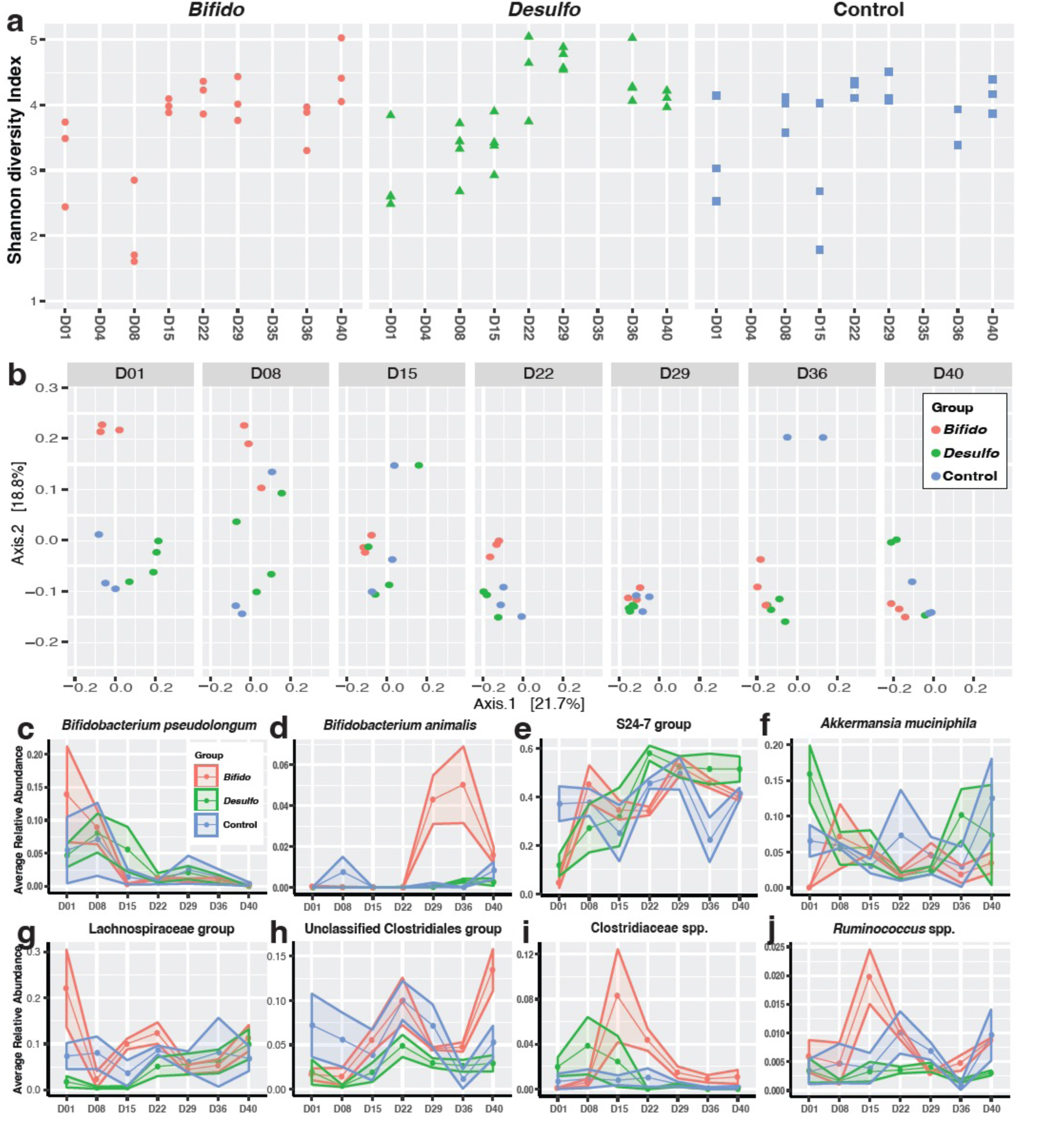
Profound early influence of SBT on gut microbiota with subsequent loss of colonization and distinctive gut microbiota features after 2 weeks. Mice given abx (6 days), then gavage (WT stool/*Bifido* MD*/Desulfo*) on the day of heart transplant, followed by tacrolimus for 40 days post-transplant (3 mice/group). Stool pellets collected longitudinally through day 40. (a) The α diversity (within-sample diversity) was estimated using the Shannon diversity index. (b) β-Diversity (between-sample diversity) was determined through multidimensional scaling plots using Jensen–Shannon divergence distance to visualize the level of divergence at the different sampling times.

Further taxonomic analysis revealed several major species responsible for the altered gut microbiota composition and structure following SBT **(Fig. 1c-j)**. The change in relative abundance of all taxonomic groups is shown in **Suppl. Fig. S1**. Most outstandingly, we observed *Bifidobacterium*, *Bacteroides*, and *Clostridiales* were among the groups most responsive to SBT. Notably, *B. pseudolongum* was most abundant after SBT with *Bifido*, although it was present at lower levels in *Desulfo* and control groups, but sharply decreased within 2 weeks in the *Bifido* treated group **(Fig. 1c)**. This decrease in *B. pseudolongum* was accompanied by a steep increase in *Bifidobacterium animalis*, a commensal that cohabitates with *B. pseudolongum,* **(Fig. 1d)**. *Bacteroidales* family S24-7, a group of fermentative anaerobic commensals that predominate in murine gut microbiota^33^, sharply decreased after SBT, but was restored from week 2 onward **(Fig. 1e)**. *Akkermansia muciniphila*, a commensal known for its anti-inflammatory and mucin-degrading properties^34^, sharply increased only in response to *Bifido* gavage **(Fig. 1f)**. *A. muciniphila* reached peak abundance at week 2, followed by a stable, reduced level through week 6. In our previous study^31^, *A. muciniphila* was enriched in the gut microbiota in *Bifido* MD gavaged mice, despite its overall low abundance. *Lachnospiraceae* and *Ruminococcus*, obligate anaerobes within *Clostridiales* with diverse fermentative capabilities, were highly variable following SBT **(Fig. 1g-j)**. Overall, these results indicate major and dynamic changes in gut microbiota composition and structure over time following SBT. We hypothesize that the systemic effects of SBT take place early and attenuate over time, as the immune system re-equilibrates and as the microbiota reverts over time.

### Early changes in intestinal and LN immune compartment structures due to *Bifido* and *Desulfo*

To investigate early immune changes triggered by individual microbiota, mice were next treated with abx (6 days), and SBT (*Bifido* MD, or *Desulfo)* or PBS control, on day 0. Tissues were obtained for analysis at day 3. There were relatively few changes in cellularity noted by flow cytometry. F4/80+ MΦ were decreased in mesenteric LN (mLN) with *Bifido* MD compared to *Desulfo* and control while CD11c+ DCs were decreased in the peripheral LN (pLN) of all treatment groups compared to control. **(Suppl. Fig. S2a)**. By immunohistochemistry (IHC), in the mLN, Foxp3+ Tregs were increased in CR with *Bifido* and *Desulfo* treatments. F4/80+ MΦ were decreased in CR and the high endothelial venules (HEV) by *Bifido* MD and *Desulfo* **(Fig. 2a,b, Suppl. Fig. S2b-e)**. In pLN, CD11c+ DCs and F4/80+ MΦ were increased compared to control with *Bifido MD* in CR **(Fig. 2c, d)**. There were no changes in pLN laminin α4:α5 ratios **(Fig. 2c)**. In the intestine, *Desulfo* treated mice showed increased Foxp3+ Tregs compared to *Bifido* and control **(Fig 2e, f)**. *Bifido* MD mice had decreased CD11c+ DCs compared to *Desulfo* and control. F4/80+ MΦ were decreased in all treatment groups compared to control **(Fig. 2e)**. *Desulfo* and *Bifido* overall resulted in a mixture of pro-and anti-inflammatory changes in immune cell distribution within secondary lymphoid organs and intestine. However, neither affected LN architecture at this early time point.

**Figure 2.**
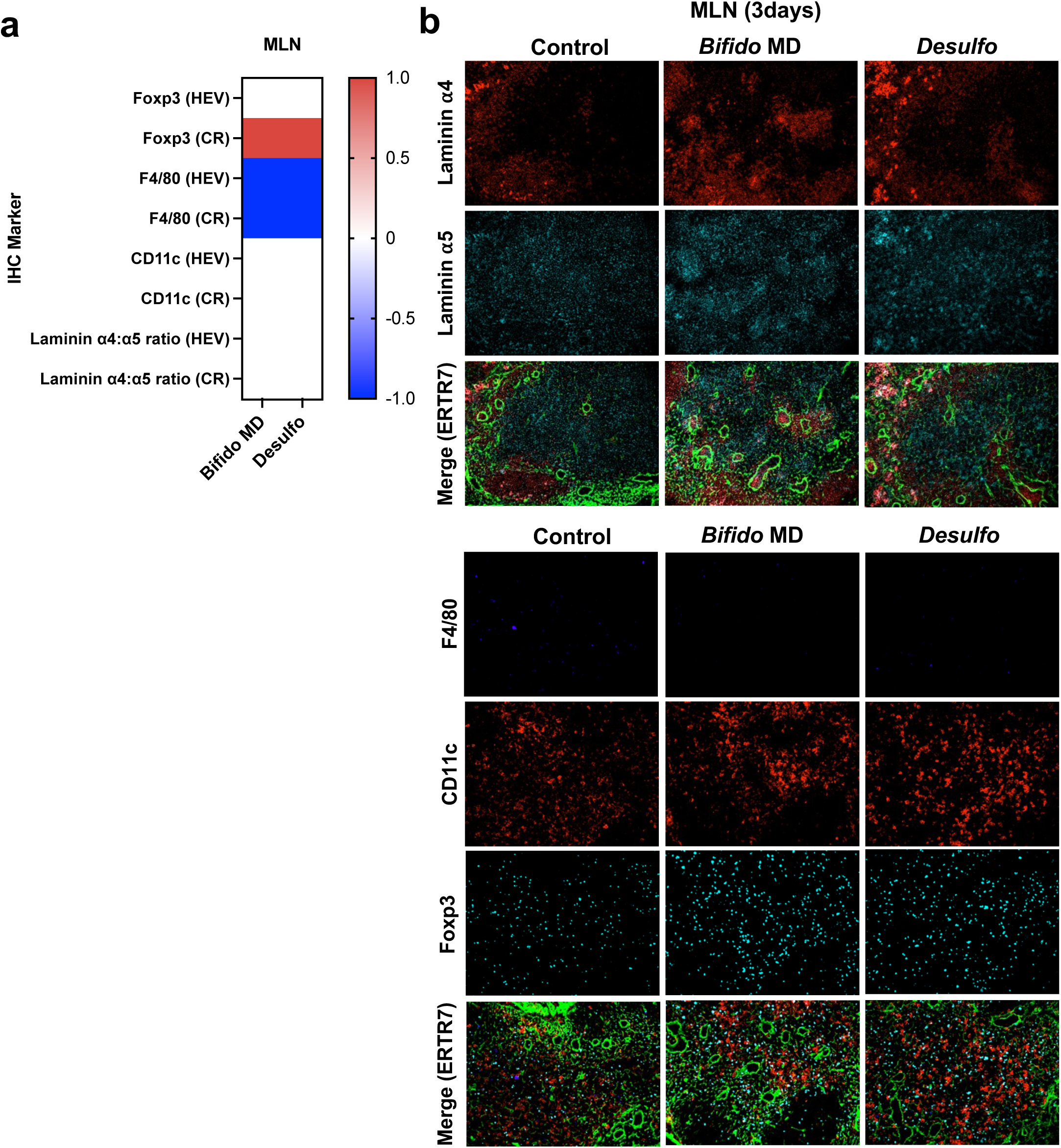
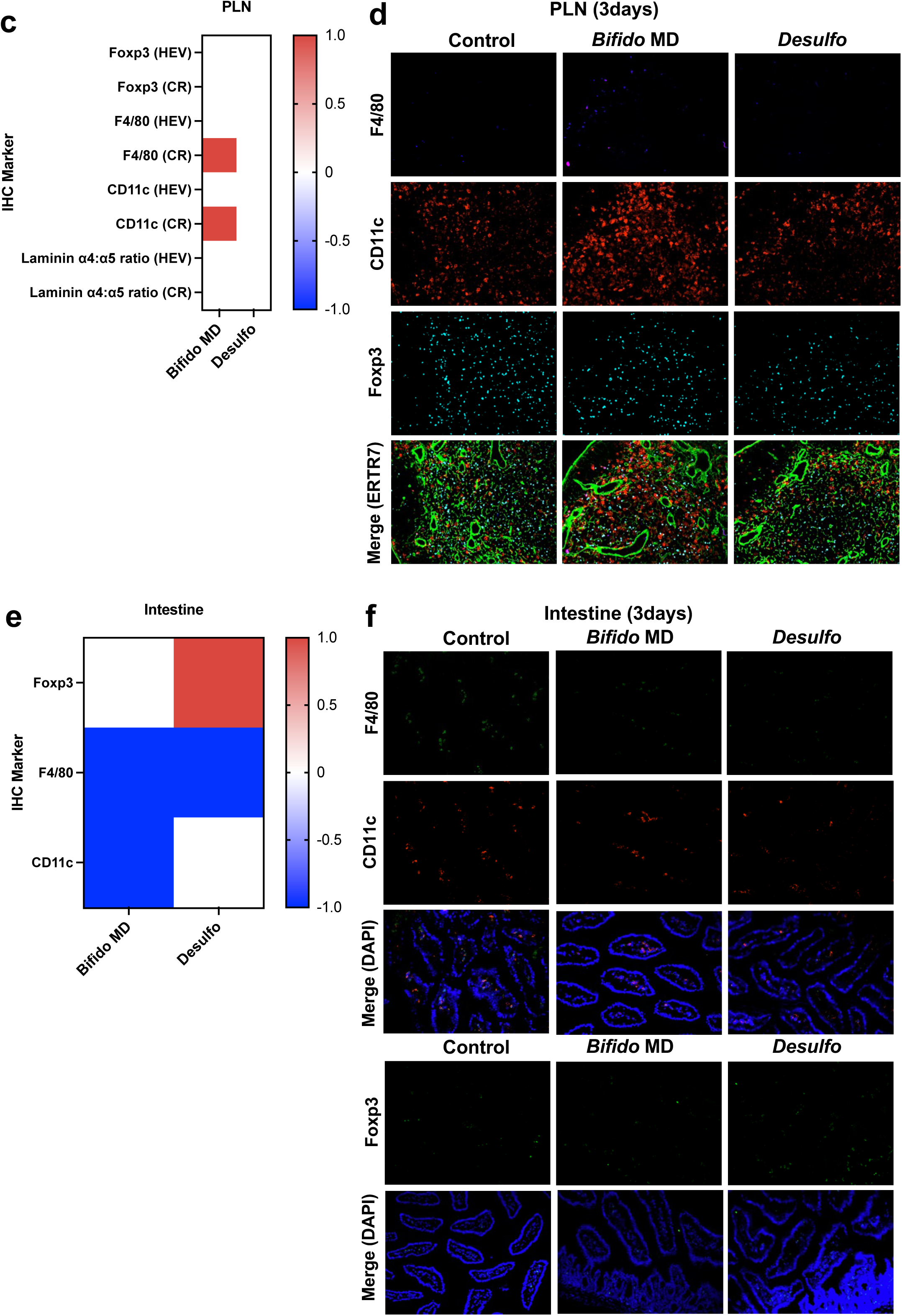
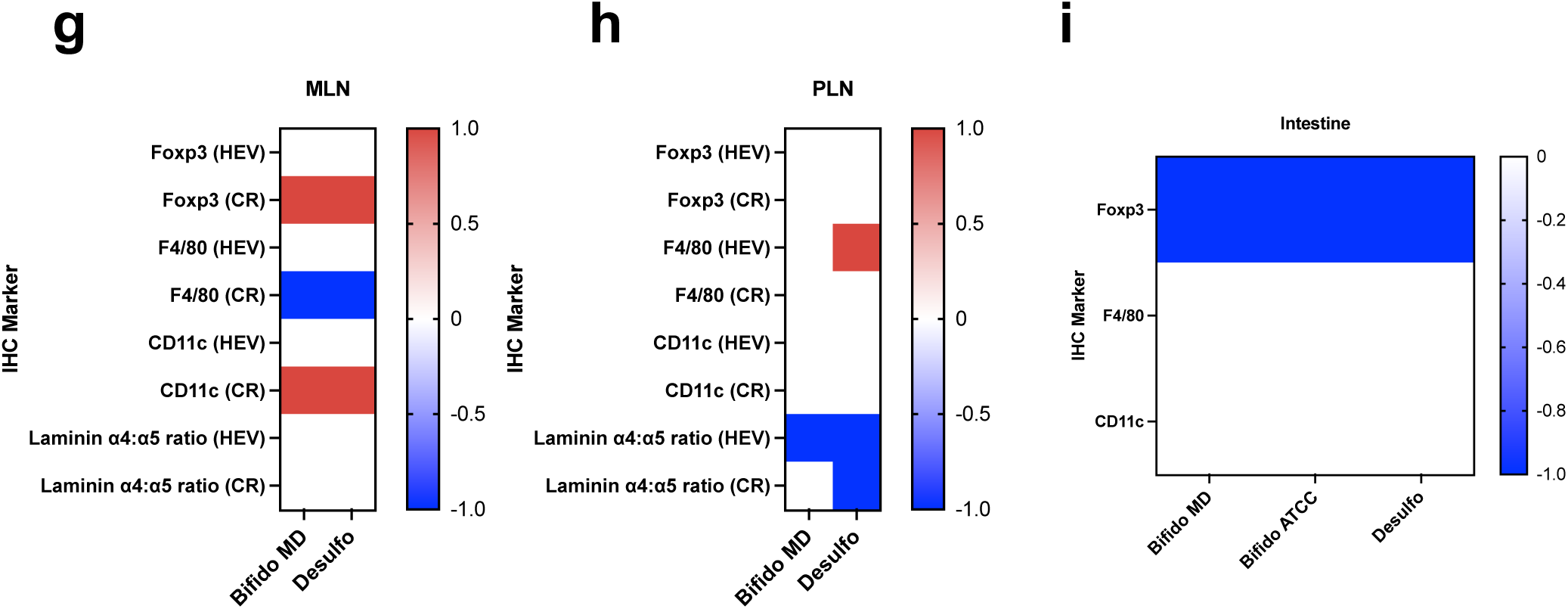
Early changes in LN architecture and immune cell distribution due to SBT. Mice given abx (6 days), then SBT (PBS/*Bifido* MD/*Desulfo*). After 3 days, LN, and intestinal segments harvested. Qualitative heat maps of IHC marker changes relative to control (red=increased; blue=decreased; white=unchanged; figure generation detailed in *methods* section) of **(a)** mLN. Representative IHC images for **(b)** mLN Laminin *α*4, *α*5, F4/80, CD11c, Foxp3, and ERTR7 (merge), 20x. Qualitative heat maps of IHC marker changes relative to control of **(c)** pLN and representative IHC images for **(d)** pLN F4/80, CD11c, and DAPI (merge). Qualitative heat maps of IHC marker changes relative to control of **(e)** intestine and representative IHC images for **(f)** intestine F4/80, CD11c, Foxp3, and DAPI (merge). Mice treated with abx (6 days), then SBT (*PBS, Bifido MD,* or *Desulfo*), followed by tacrolimus for 2 days. After 3 days, LN and intestinal segments harvested. Representative qualitative heat maps of IHC marker changes relative to control of **(g)** mLN, **(h)** pLN, and **(i)** intestine. Data representative of 2 separate experiments, 3 mice/group, at least 2 mLN, pLN, or segments of intestine/mouse, 3 sections/tissue/staining panel, and 3–5 fields imaged/section, i.e., 20-30 total microscopy fields/condition over 2 experiments. Ordinary one-way ANOVA with Tukey’s multiple comparisons test. * p < 0.05; ** p < 0.01, *** p < 0.001, **** p < 0.0001.

Incorporating the clinically relevant immunosuppression tacrolimus eliminated the minor changes observed by flow cytometry in mLN, pLN, and spleen after 3 days **(Suppl. Fig. S3a-b)**. While the absolute numbers and ratios of cell content did not change as assessed by flow cytometry in mLN, the microanatomic distribution of these cells around the CR did change. There were also increased mLN CR laminin α4:α5 ratios with *Bifido* MD, **(Fig. 2g, Suppl. Fig. S3c).** In pLN HEV and CR, there were decreased laminin α4:α5 ratios with *Desulfo* **(Fig. 2h, Suppl. S3d)**. In the pLN HEV, *Bifido* MD treatment resulted in decreased laminin ratios. IHC of intestinal segments showed a decrease in Foxp3+ Tregs cells in all treatment groups **(Fig. 2i, Suppl. Fig. S3e)** suggesting that tacrolimus inhibited early intestinal changes from abx and SBT. Overall, *Bifido* resulted in anti-inflammatory changes in the mLN with increased laminin ratios while causing simultaneous pro-inflammatory changes in pLN and intestine with decreased laminin ratios and Foxp3+ Tregs, respectively. *Desulfo* displayed a largely pro-inflammatory phenotype with decreased pLN laminin ratios and decreased Foxp3+ Tregs in the intestine despite increased mLN Foxp3+ Tregs.

We next studied the capability of tacrolimus to modulate the alloimmune response to abx and SBT in transplanted recipients. Mice were treated with abx, SBT (*Bifido* MD, *Desulfo*, or PBS control), cardiac transplantation, and tacrolimus. Tacrolimus was continued daily for 7 days, and recipients euthanized on day 8 after transplant. In the mLN and pLN, there were no significant changes in Foxp3+ Tregs or innate immune populations between treatment groups. *Desulfo* resulted in increased laminin α4:α5 ratios in the mLN CR only **(Fig. 3a, Suppl. Fig. S4a)**. In the intestine, *Desulfo* treated mice had decreased CD11c+ DCs compared to control **(Fig. 3b)***. Bifido* MD resulted in decreased intestine Foxp3+ Treg. Overall, SBT resulted in few, relatively small differences in immune cell distribution within LN and intestine at this early time point.

**Figure 3.**
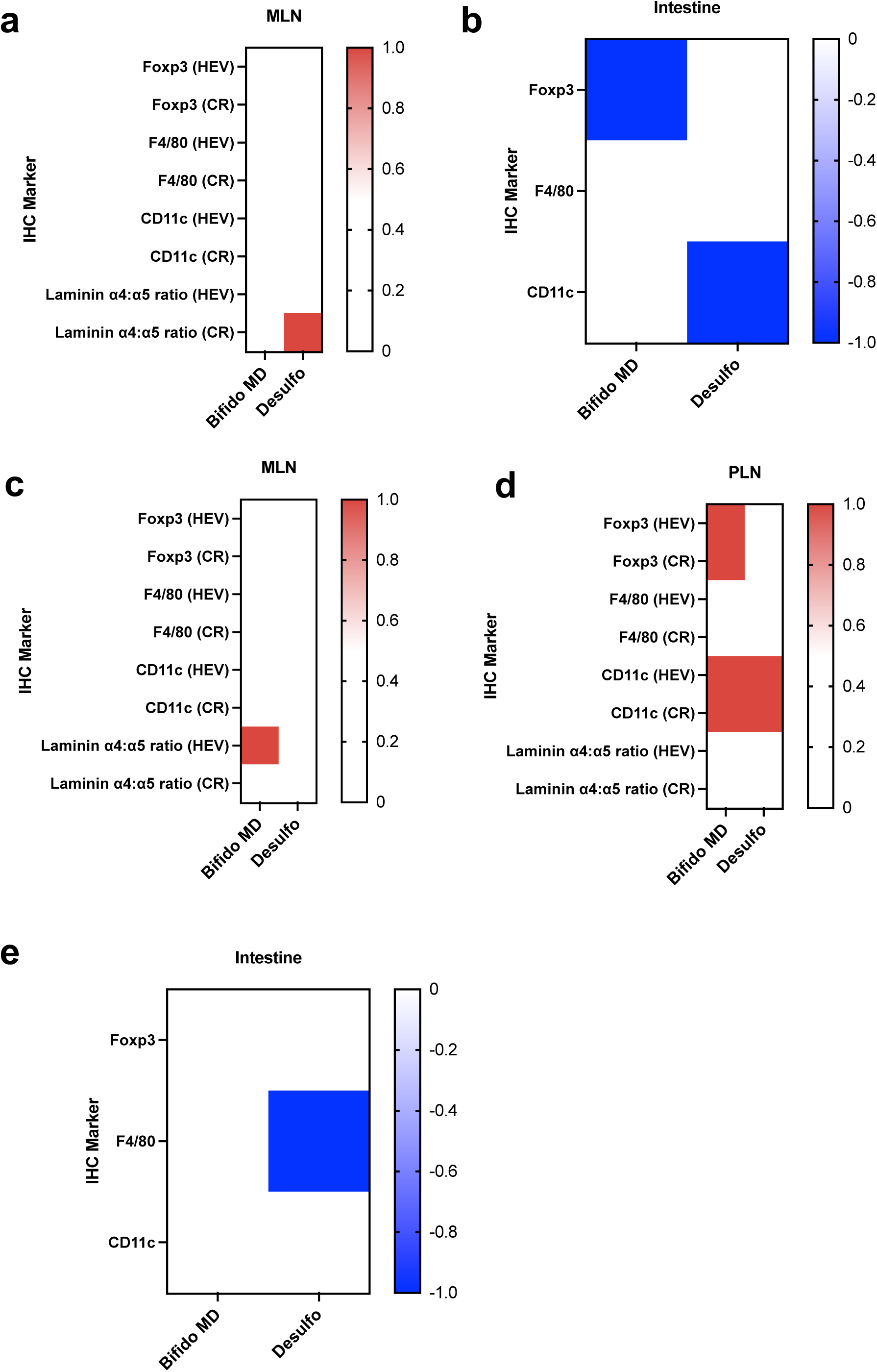
Changes in LN structure and immune cell distribution due to abx, SBT, heart transplant, and tacrolimus at 7 and 14 days. **(a)** Mice given abx (6 days), then SBT (PBS/*Bifido* MD/*Desulfo*) on the day of heart transplant with injection of TEa T cells (2 x 10^5^ cells i.v.), followed by tacrolimus for 7 days. Representative qualitative heat maps of IHC marker changes relative to control of **(a)** mLN and **(b)** intestine. **(c-e)** Mice given abx (6 days), then SBT (PBS/*Bifido*/*Desulfo*) on the day of heart transplant with injection of TEa T cells (2 x 105 cells i.v.), followed by tacrolimus for 14 days. Representative qualitative heat maps of IHC marker changes relative to control of **(c)** mLN, **(d)** pLN, and **(e)** intestine. Data representative of 2 separate experiments with 3-5 mice/group, at least 2 mLN, pLN, or segments of intestine/mouse, 3 sections/tissue/staining panel, and 3–5 fields imaged/section, i.e., 20-30 total microscopy fields/condition over 2 experiments. Ordinary one-way ANOVA with Tukey’s multiple comparisons test. * p < 0.05; ** p < 0.01, *** p < 0.001, **** p < 0.0001.

We further assessed this immunomodulatory effect after 14 days. By IHC, there were no changes in Foxp3+ Tregs and innate immune cells in mLN HEV or CR **(Fig. 3c, Suppl. Fig. S4b-e)**. However, *Bifido* MD resulted in increased laminin α4:α5 ratios around mLN HEV compared to control and *Desulfo* (**Fig. 3c)**. In pLN, *Bifido* treatment resulted in increased Foxp3+ Tregs in HEV compared to control and CR compared to both *Desulfo* and control. *Desulfo* had increased CD11c+ DCs in CR and HEV compared to control (**Fig. 3d)**. There were no differences in the ratios of laminin expression with *Bifido* MD treatment in pLN. In the intestine, *Desulfo* treated mice had decreased F4/80 MΦ compared to control **(Fig. 3e)**. Overall, *Bifido* MD resulted in pro-tolerogenic changes in pLN with increased Foxp3+ Tregs and mLN with increased laminin ratios, whereas *Desulfo* appeared to have no effect at day 14.

### *Bifido* and *Desulfo* induce early changes in T cell responses

Given the changes in immune cell content and LN architecture early on after treatment with specific microbiota strains, we sought to characterize its impact on antigen specific T cell trafficking and differentiation. Mice were treated with abx, SBT (*Bifido* MD, *Desulfo*, or PBS control), cardiac transplantation, and tacrolimus. Mice also received adoptive transfer of alloantigen specific, T cell receptor transgenic, CD4+ TEa T cells (2 x 10^5^) on the day of transplant. Recipients were euthanized on day 8. After 7 days, the percentages of transferred TEa cells in mLN, pLN, or spleen did not differ between treatment groups, although there was a non-statistically significant trend toward increased TEa percentages in *Desulfo* treated animals, possibly reflecting increased proliferation. TEa cells differentiated into similar percentages of Foxp3+ Tregs in all groups. *Bifido* treated animals demonstrated an increased percentage of naïve (CD4+CD44-CD62L+CD69-) TEa T cells (**Fig. 4a**). There was an increased percentage of effector memory (CD4+CD44+CD62L-CD69-) TEa T cells in *Desulfo* treated spleen and pLNs compared to the *Bifido*, with a similar trend in mLN (**Fig. 4a**). These findings reflect a pro-inflammatory, activating effect of *Desulfo* while *Bifido* demonstrates a pro-homeostatic effect on naïve CD4+ T cell populations.

**Figure 4.**
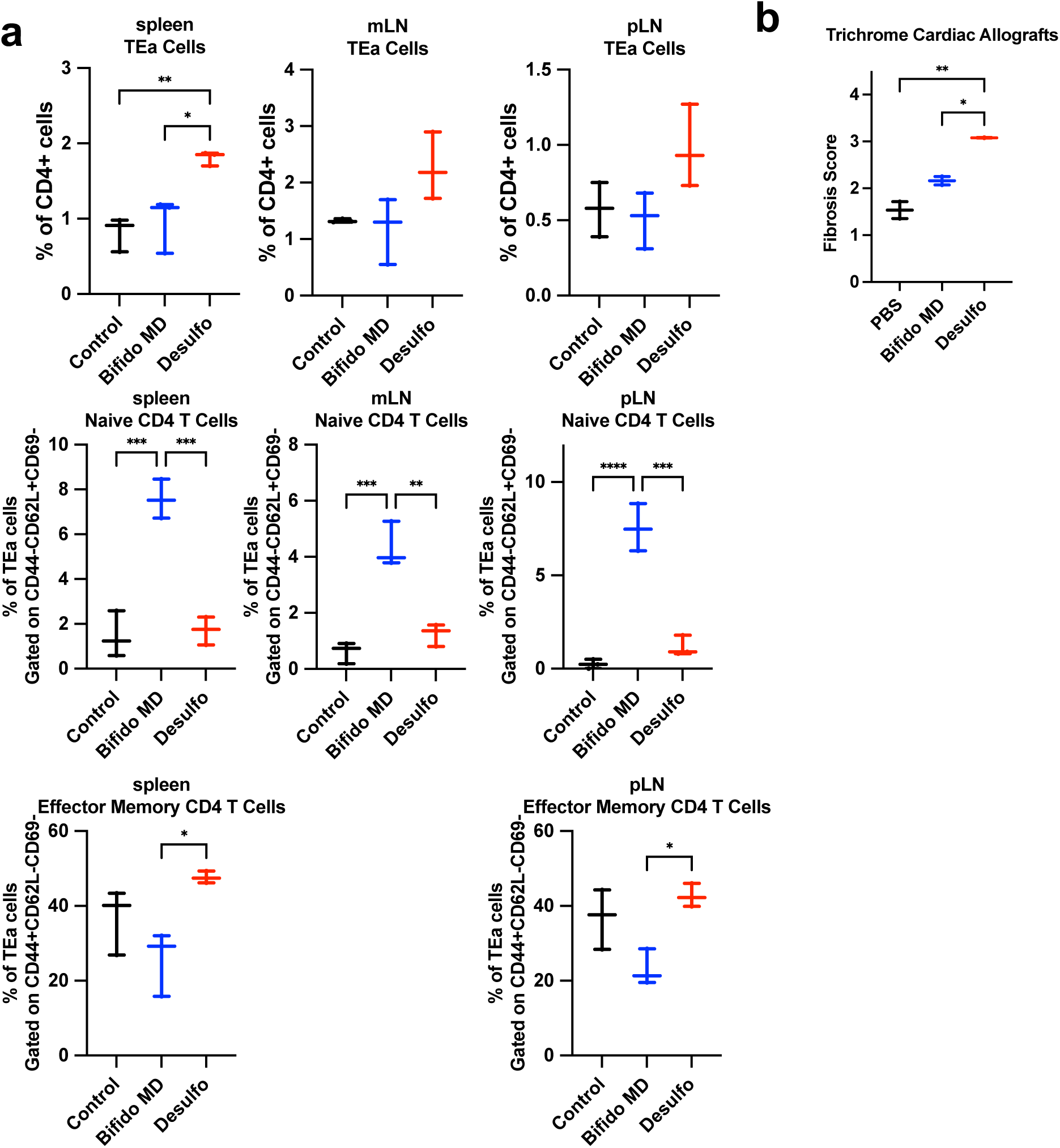
Early changes in T cell and allograft responses due to abx, SBT, heart transplant, and tacrolimus at 7 and 14 days. Mice given abx (6 days), then SBT (PBS/*Bifido* MD/*Desulfo*)on the day of heart transplant with injection of TEa T cells (2 x 10^5^ cells i.v.), followed by tacrolimus for 7 days. **(a)** Flow cytometry of mLN, pLN, and spleen for CD4, TEa TCR, Foxp3, CD44, CD62L, and CD69. **(b)** Mice given abx (6 days), then SBT (PBS/*Bifido/Desulfo*) on the day of heart transplant with injection of TEa T cells (2 x 105 cells i.v.), followed by tacrolimus for 14 days. Cardiac allograft fibrosis by trichrome stain. Data representative of 2 separate experiments with 3-5 mice/treatment group/experiment. Ordinary one-way ANOVA with Tukey’s multiple comparisons test. * p < 0.05; ** p < 0.01, *** p < 0.001, **** p < 0.0001.

Graft inflammation and fibrosis were very low and similar across the 3 study groups, as expected for this very early post-transplant time point (**Suppl. Fig. S5a**). However, allografts had greater fibrosis after 14 days in *Desulfo* treated animals compared to *Bifido* or control **(Fig. 4b)** consistent with our previous findings^30^. Graft inflammation though remained similar across the 3 study groups **(Suppl. Fig. S5a, b)**. The gavage treatments did not affect serum tacrolimus levels across study groups (**Suppl. Fig. S5c**).

### *Bifido* and *Desulfo* result in changes in long term graft health and LN laminin **α**4:**α**5 ratios

Having shown that SBT induces immune alterations within a relatively short time period, while microbiome data revealed short-lived stability of *Bifido* colonization, we sought to test whether these early changes left long lasting effects on allograft tolerance and survival. Mice were treated with abx, gavaged with WT mouse whole stool samples (control) or SBT (*Bifido* or *Desulfo*). Cardiac transplants were then performed and followed by daily low dose tacrolimus immunosuppression. Allografts were assessed for inflammation and fibrosis 40 days after transplant along with LNs for architectural changes characterized by the ratio of laminin α4:α5 and the presence of Foxp3+ Tregs in the CR and around the HEV.

Mice that received porcine-derived *B. pseudolongum* ATCC27774 (*Bifido* ATCC) SBT had lower cardiac allograft inflammation scores than mice gavaged with *Desulfo* or treated with control gavage (**Fig. 5a**), consistent with our previous results^30^. Histology revealed more CD4+ helper and CD8+ cytotoxic T lymphocytes in the grafts of *Desulfo* treated mice (**Fig. 5b**). The LN laminin α4:α5 ratio in the CR and around the HEVs, the regions where Foxp3+ Tregs are induced and activated, were significantly higher in *Bifido* ATCC gavaged mice compared to WT stool control mice, suggestive of a more suppressive environment (**Fig. 5c**). Of note, our group’s original studies utilized commercially available, porcine-derived *Bifido* ATCC^30^ prior to isolating and growing monocultures of murine-derived *Bifido* MD^35^. In comparing *Bifido* ATCC and MD immunologic effects over longer time periods of 40 and 60 days, we found their immunologic effects on allografts and LN to be highly similar **(Fig. 5d)**, despite some short-term strain-specific *in vitro* differences^31^.

**Figure 5.**
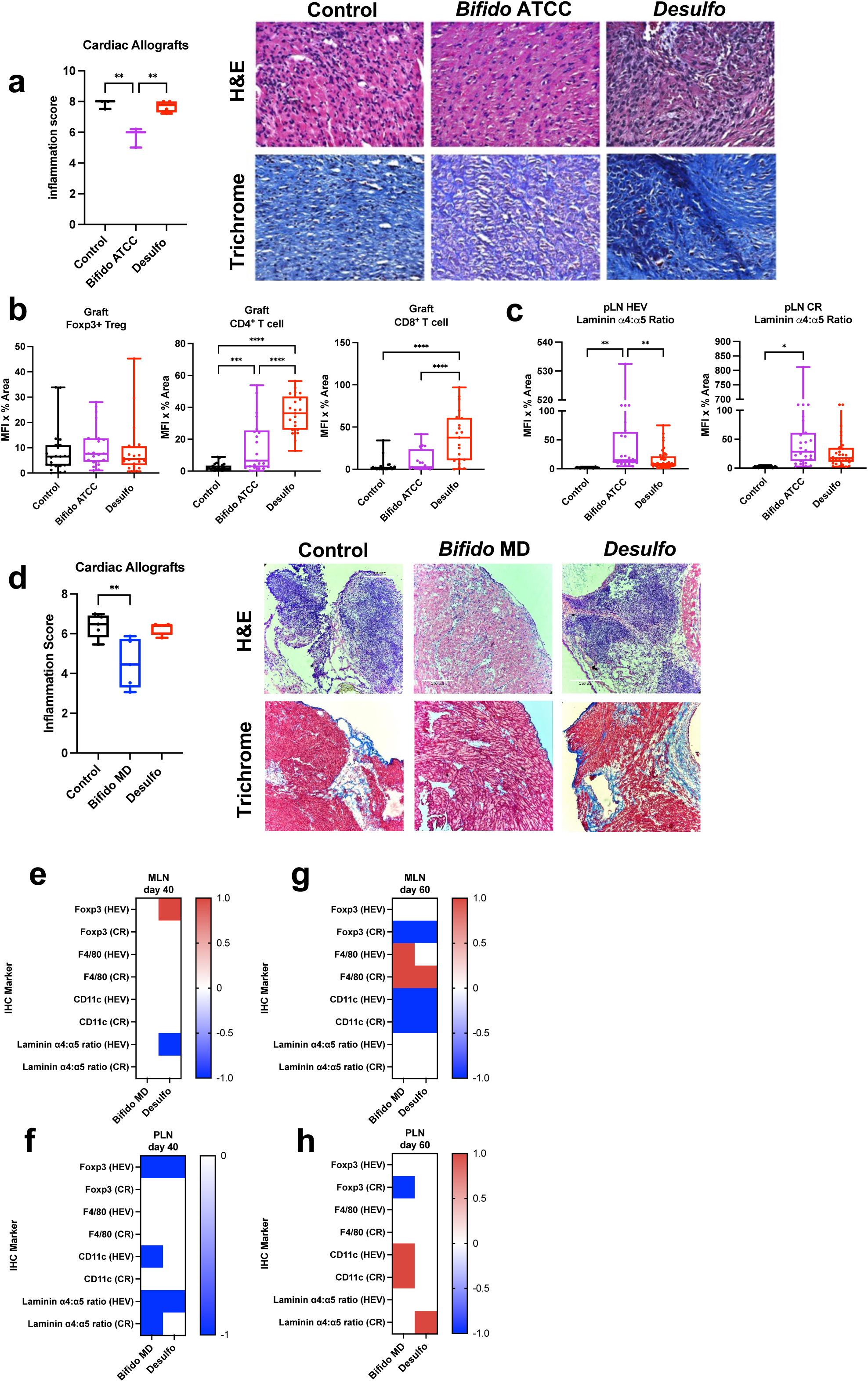
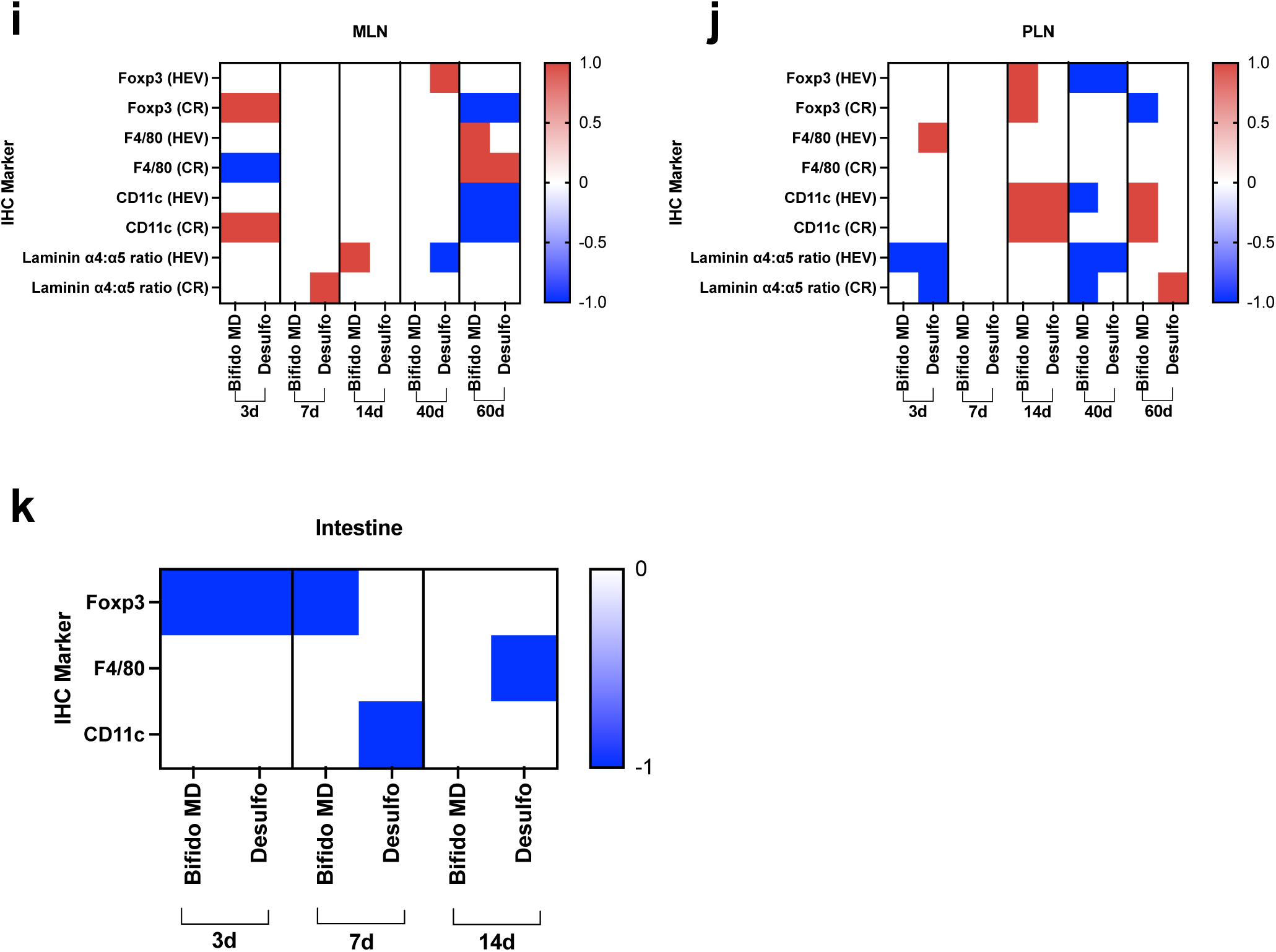
Effect of *Bifido* and *Desulfo* compared to normal SBT on cardiac graft inflammation and LN remodeling. Mice given abx (6 days), then gavage (WT stool/*Bifido ATCC/Desulfo*) on the day of heart transplant, followed by tacrolimus for 40 days. **(a)** Graft inflammation scores with representative histology images (H&E and trichome stains, 20x). **(b)** Foxp3+ Tregs, CD4, and CD8 T cells in the cardiac grafts. **(c)** Laminin *α*4, *α*5, and ratios in pLNs. Mice given abx (6 days), SBT (PBS/*Bifido* MD/*Desulfo*) on the day of heart transplant, followed by tacrolimus for 40 days. **(d)** Graft inflammation scores with representative histology images (H&E and trichome stains, 20x). Representative qualitative heat maps of IHC marker changes relative to control of **(e)** mLN and **(f)** pLN. Mice given abx (6 days), SBT (PBS/*Bifido* MD/*Desulfo*) on the day of heart transplant, followed by tacrolimus for 60 days. Representative qualitative heat maps of IHC marker changes relative to control of **(g)** mLN and **(h)** pLN. Representative qualitative heat maps of IHC markers were pooled for 3, 7, 14, 40, and 60 day experiments for **(i)** mLN, **(j)** pLN, as well as 3, 7, and 14 day experiments for **(k)** intestine. 40-day data representative of 3 separate experiments, 60-day data represent of single experiment. Three mice/group, at least 2 mLN, pLN, or segments of intestine/mouse, 3 sections/tissue/staining panel, and 3–5 fields imaged/section, i.e., 20-30 total microscopy fields/condition. Ordinary one-way ANOVA with Tukey’s multiple comparisons test. * p < 0.05; ** p < 0.01, *** p < 0.001, **** p < 0.0001.

Since *Bifido* MD is murine-tropic with improved anti-inflammatory capacity *in vitro*^31^, subsequent transplant experiments were carried out using the *Bifido* MD strain. We compared *Bifido* MD SBT to *Desulfo* or control PBS gavage in animals receiving allogeneic cardiac transplants. There were no episodes of early rejection and animals were sacrificed at day 40. Similar to *Bifido* ATCC, *Bifido* MD treatment resulted in decreased allograft inflammation based on histology compared to control **(Fig. 5d)**. By flow cytometry, there were minimal changes between SBT groups for CD4+ or CD8+ T cells, Foxp3+ Tregs, CD11c+ DCs, or F4/80+ MΦ populations in mLN, pLN, or spleen (**data not shown)**. By IHC, there were increased Foxp3+ Tregs around mLN HEV of *Desulfo* treated mice **(Fig. 5e, Suppl. Fig. S6a-d)**. *Bifido* MD and *Desulfo* treated mice had decreased laminin α4:α5 ratios around the mLN HEV **(Fig. 5e)**. Notably, *Desulfo* treatment resulted in decreased laminin ratios in both HEV and CR compared to *Bifido* MD, consistent with a more inflammatory immune environment. In pLN, *Bifido* MD and *Desulfo* treated mice had decreased Foxp3+ Tregs around HEVs along with decreased laminin ratios around HEV (**Fig. 5f)**. Overall, *Desulfo* had mixed effects in mLN with increased anti-inflammatory Foxp3+ Treg as well as decreased laminin ratios, while in the pLN, *Desulfo* was more pro-inflammatory with decreased Tregs and laminin ratios. *Bifido* MD exerted pro-inflammatory changes as well with decreased pLN Foxp3+ Tregs and decreased pLN laminin ratios.

We next investigated whether these changes persisted 60 days after transplant and SBT.

By flow cytometry, splenic CD4+ and CD8+ T cells were decreased in *Bifido* MD treated mice compared to control. In the pLN of *Bifido* MD treated animals, F4/80+ MΦ were decreased compared to control **(Suppl. Fig. S7a)**. By IHC, *Bifido* MD and *Desulfo* treated mice had decreased Foxp3+ Tregs in the mLN CR **(Fig. 5g, Suppl. Fig. S7b-e)**. In pLN, *Bifido* MD treated mice had decreased Foxp3+ Tregs in the CR **(Fig. 5h)**. In the pLN CR, *Desulfo* treated mice showed increased laminin ratios compared to both control and *Bifido* **(Fig. 5h)**. Overall, on day 60, *Bifido* MD mice showed decreased splenic T cells and increased B cells along with decreased pLN F4/80+ MΦ by flow cytometry. By IHC, *Bifido* MD showed pro-inflammatory effects with decreased Foxp3+ Tregs in pLN and decreased laminin ratios in mLN, similar to day 40. *Desulfo* resulted in mixed effects with pro-inflammatory mLN changes with decreased Foxp3+ Tregs in mLN and pLN as well as increased anti-inflammatory laminin ratios in pLN, similar to its effects at day 40. Single dose administration of *Bifido MD* and *Desulfo* each elicited significant differences in LN architecture and immune cell homing compared to one another. However, the changes caused by SBT and tacrolimus (3 days) or by SBT, tacrolimus, and heart transplant (7, 14, 40, and 60 days) in LNs were mostly transient when comparing *Bifido* MD and *Desulfo* treatments **(Fig. 5i-k)**. More notable and persistent differences were observed in allograft histology (14 and 40 days) and alloantigen T cell homing and differentiation (7 days), suggesting that while secondary lymphoid organs return to a homeostatic baseline relatively quickly after SBT, immunomodulatory effects on the allograft are more long-lived.

### *Bifido* and *Desulfo* impact the activation programming of DC and MΦ *in vitro*

The gut is the largest immune organ and functions as a physical barrier against luminal microbes while also serving as a critical interface between commensal microbiota and innate immune cell populations^36,37^. We next investigated the direct interaction between myeloid cells and members of gut microbiota by incubating primary bone marrow derived DCs (BMDCs) and peritoneal MΦ with UV-killed *Bifido* MD or *Desulfo*. After 24 hours, supernatants were assessed for IL-6, IL-10, and TNFα by ELISA; and cells analyzed by flow cytometry for activation markers. BMDCs displayed increased MHC II in response to *Desulfo*. *Bifido* MD and *Desulfo* all increased CD40 expression, with *Desulfo* stimulating the greatest increase. Both *Bifido* MD and *Desulfo* increased CD80 expression, whereas only *Desulfo* increased CD86 **(Fig. 6a)**. The percentage of M1 MΦ (F4/80+ MHC II+) was not affected by treatment **(Fig. 6b)**. *Bifido* MD treatment led to an increased percentage of M2 MΦ (F4/80+ CD206+) with *Desulfo* inducing a greater percentage of M2 than *Bifido* MD. CD40 expression increased after either bacterial treatment among the M1 subset (**Fig. 6b)**. These data demonstrate that gut microbiota directly activate antigen presenting cells (APCs), with each strain triggering a unique activation pattern.

**Figure 6.**
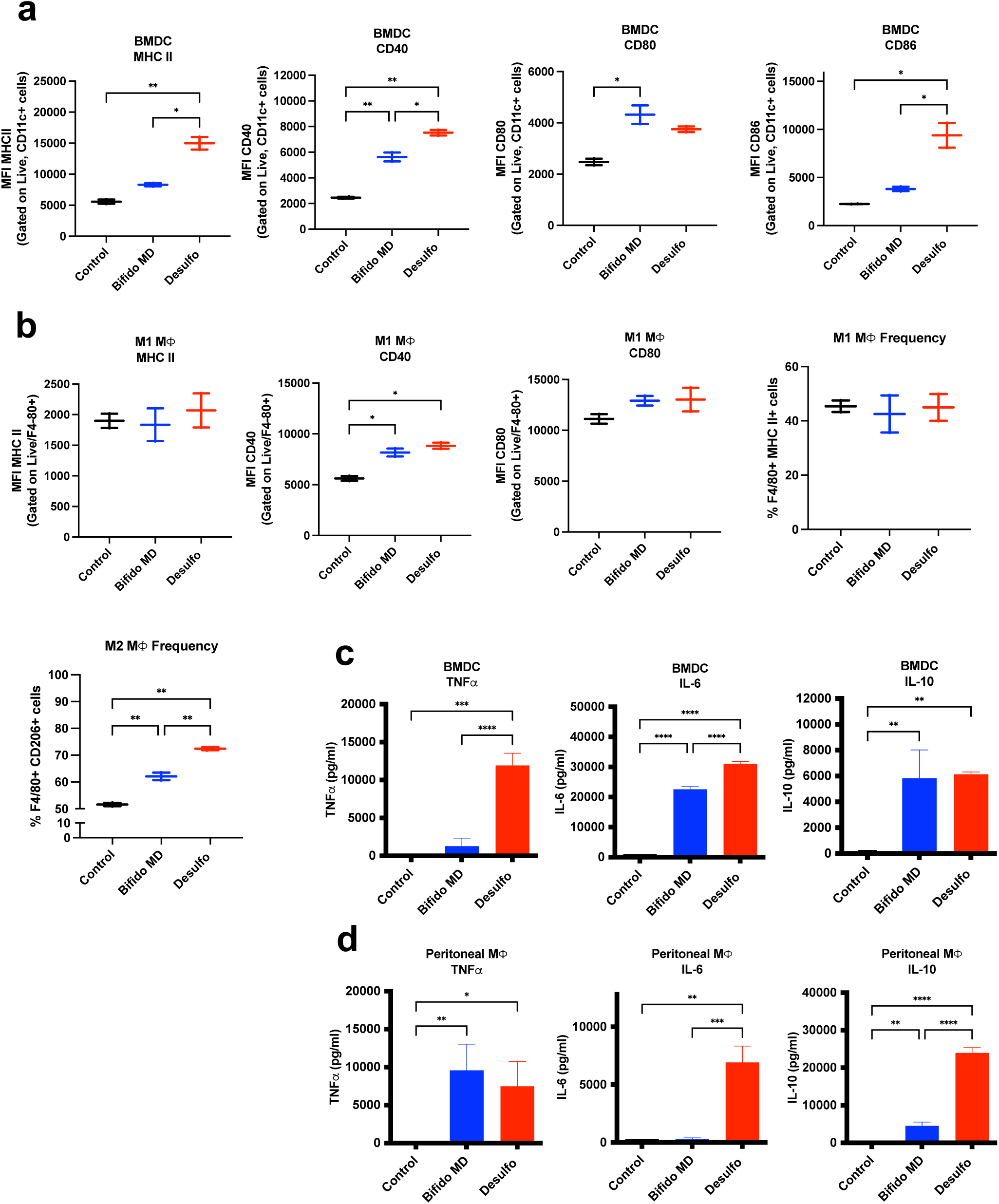
Microbiota directly modulate activity of DC and M<Φ *in vitro*. BMDCs and freshly isolated peritoneal M<Φ stimulated with UV-killed *Bifido* MD, *Desulfo*, or media alone. After 24 hours **(a)** BMDC and **(b)** M<Φ were analyzed by flow cytometry for MHC II, CD40, CD80, and CD86. **(c)** BMDC and **(d)** M<Φ culture supernatants assessed for IL-6, IL-10 and TNFα by ELISA. M<Φ flow cytometry data representative of 2 separate experiments and DC flow cytometry data representative of single experiment, 2 wells/culture condition. ELISA data representative of 2 separate experiments, 2-3 wells/culture condition, 2 technical replicates/well (supernatants from each well split and analyzed in duplicate), i.e., 4 wells/condition/experiment, 10 total wells/condition over 2 experiments. Ordinary one-way ANOVA with Tukey’s multiple comparisons test. * p < 0.05; ** p < 0.01, *** p < 0.001, **** p < 0.0001.

In BMDCs, TNFα was elevated after *Desulfo*, but not *Bifido* MD stimulation **(Fig. 6c)**. IL- 6 was elevated in all groups with progressively higher levels from *Desulfo* compared to *Bifido* MD. IL-10 was elevated in all treatment groups **(Fig. 6c)**. Overall, BMDC responses to *Bifido* were relatively anti-inflammatory (decreased TNFα and IL-6; increased IL-10) compared to *Desulfo.* In peritoneal MΦ, TNFα was elevated by both *Desulfo* and *Bifido* MD **(Fig. 6d)**. *Desulfo* increased IL-6. *Bifido* MD increased IL-10, while *Desulfo* resulted in even higher levels **(Fig. 6d).** Overall, *Desulfo* induced a pro-inflammatory phenotype in MΦ with elevated IL-6 and TNFα and stimulated more IL-10 compared to *Bifido*. Notably, IL-10 is not exclusively immunosuppressive, but can augment the differentiation of B and CD8+ T cells while also accelerating heart allograft rejection^38^.

## DISCUSSION

This study recapitulated our initial findings of the deterministic role that single bacterial strains play in pro-and anti-inflammatory host immune modulation^30^. Not only were specific bacterial strains able to skew alloimmune responses toward tolerance or rejection in the short term, but we also showed that abx, immunosuppression, and transplantation each modified this process. Most outstandingly, we demonstrated the time-dependent effect of pro-tolerogenic and pro-inflammatory bacterial strains (*Bifido* and *Desulfo*, respectively) on immunological events and graft outcomes. In particular, we observed short term, transient changes in secondary lymphoid organs and intestine as well as in the gut microbiome with each bacterial treatment, which resolved over time; however, these initial changes still resulted in major differences in allograft inflammation and fibrosis over time. We also showed that each bacteria acted directly on APCs *in vitro* yielding specific effects on surface co-stimulation markers and cytokine production.

We previously showed that *Bifido* and *Desulfo* gavage were able to exert pro-tolerant and pro-inflammatory LN architecture effects after 14 days, respectively^30^. In this study, we expanded on these findings to show that *Bifido* and *Desulfo* affected LN immune cell content and distribution as early as 3 days following gavage. Increased allograft fibrosis was observed with pro-inflammatory *Desulfo* after 14 days and decreased inflammation in allografts in mice treated with pro-tolerant *Bifido* after 40 days. However, the majority of bacterial effects on secondary lymphoid organs were most pronounced at early time points, becoming obscured over time, as evidenced by both pro and anti-inflammatory changes in LN architecture and immune cell content and distribution beyond 14 days of treatment. We speculate that these kinetic changes may be the result of complex interactions among the multiple treatments each mouse received. First, long-term immunosuppression can lead to dysbiosis in human transplant recipients^39^. Preliminary work by our group has confirmed that treatment with immunosuppression or abx alone significantly alters the gut microbiome and metabolome after 3 days, suggesting the importance of microbial metabolites as immunologic mediators^26^. Second, abx can significantly impede the colonization of specific bacterial strains due to increased colonization by antimicrobial drug resistance strains, resulting in blooms of pathogenic bacteria^42,43^. Third, the availability of nutrient niches within the host microbiota strongly influences the colonizing potential and stability of individual administered bacteria strains^43^. These results confirmed our earlier findings that *Bifido* exerts pro-tolerant changes and *Desulfo* exerts pro-inflammatory LN changes early on, as well as differential effects on allogeneic T cell homing and differentiation, culminating in altered degrees of alloimmunity. They also suggest that LN architecture and lymphocyte content changes are dynamic over longer treatment periods, despite appreciable long term changes to allograft histology. These observations strongly suggest that immune analyses require a detailed appreciation of several major variables and an assessment of both short and long-term changes. For microbiota/probiotic-based interventions, longitudinal monitoring of bacteria colonization and consideration of repeated administration may be critical to ensure a long term biotherapeutic effect^39,44^.

We previously showed that *Bifido* MD and *Bifido* ATCC elicited strain-specific differential effects on innate immune cell activation, tolerogenic LN architecture, and host transcriptome profiles^31^. In this study, murine-isolated *Bifido* MD elicited early pro-tolerant immune changes in terms of allogeneic T cell differentiation while histologic allograft changes were observed over longer periods. Overall, LN architectural changes were transient. The various taxonomic groups that were shown to respond to SBT in this study demonstrate the major involvement of endogenous gut microbiota as intermediaries between probiotic treatment and host immune responses. For example, the observed decrease in *B. pseudolongum* with concomitant increase in *Bifidobacterium animalis* following SBT, highlights “functional redundancy”, describing physiologic processes that can be carried out by different bacterial species at different times. Unlike endogenous *Bifido* MD, which was sourced from pregnant mice stool, exogenous porcine-derived *Desulfovibrio desulfuricans* ATCC 27774 did not colonize the murine gut post-gavage, yet still elicited a significant immune response and altered gut microbiota composition. The observed absence of *Desulfovibrio* may be attributed to colonization resistance, a known phenomenon whereby endogenous host microbiota intrinsically limits the introduction of exogenous microorganisms^42,45^. Despite this resistance, the *Desulfovibrio* strain’s capacity to trigger a robust immune response aligns with our study objectives. Thus, pro-and anti-inflammatory microbiota each caused significant changes to the gut microbiota in their own respective ways.

The gut microbiota is intimately involved with innate immunity through cells such as MΦ and DC, and their subsequent effects on the host adaptive immune response. Intestinal MΦ form an interdigitated network with the entire mucosal lamina propria vasculature; these connections are disrupted in the absence of microbiota, which also correlates with increased bacterial translocation across the lamina propria^46^. In addition to their well-known roles in antigen presentation, myeloid cells have major roles in regulating secondary lymphoid organ structure and function. Chyou *et al.*^47^ and Kumar et al.^48^ showed the importance of DCs in regulating the structure of LN vasculature and stromal cells in response to and resolution of immune responses. Matozaki and colleagues identified important roles for SIRPα+ DCs in determining the growth and differentiation of LN stromal fibroblastic reticular cells (FRC)^49,50^. FRCs in turn are responsible for constructing the laminin α-chain scaffold that promotes immunogenic or tolerogenic niches within the LN to regulate immune responses^51–53^. Together, these investigations suggest an essential mechanism whereby the microbiota stimulates gut myeloid responses, which in turn regulate regional or systemic responses via soluble mediators or migration to the LN. The LN responses include functional alterations in FRC and vascular stromal cells that determine microdomain structures in the LN that regulate adaptive immune responses. Future studies may assess the axis of communication between microbiota, APCs homing between gut and LN, and FRCs.

Our results emphasize that abx and immunosuppression alone and in concert modulate LN architecture and immune cell composition^26^. Within this network, SBT with *Bifido* induced an early anti-inflammatory phenotype while *Desulfo* resulted in a pro-inflammatory phenotype using both an *in vivo* transplant model and primary cell culture. This effect is likely related to the transient stability of each gavaged bacterial strain within the host microbiota, as evidenced by our longitudinal 16S RNA analysis of recipient gut microbiome following transplant. The gut microbiota can act directly on MΦ and DC, and the immunomodulatory effect of *Desulfo* is mediated at least partially through CCR2, as receptor blockade abrogated bacterial pro-inflammatory effects *in vivo* (not shown). *Bifido* and *Desulfo* likely act through several other mechanisms. Future studies are warranted to assess the stability of *Bifido* and *Desulfo* colonization as well as the nature of their interactions with specific host immune cells *in vivo* following bacterial gavage using longitudinal high resolution gut metagenome analysis, spatial transcriptomics profiling of gut mucosa and secondary lymphoid organs, and metabolomic analysis to probe bacteria-encoded metabolite mediators that exert critical immunomodulatory effects over time. In addition, the long term changes in allograft histology observed warrant deeper analysis of immunologic scarring and immune cell homing into allografts following bacterial treatment.

## MATERIALS AND METHODS

All materials and methods can be found in ***Supplemental Materials and Methods*.**

## AUTHOR CONTRIBUTIONS

JSB, BM, EFM, SJG, and VS conceived the study, designed experiments, interpreted the data and wrote the manuscript. SJG, AK, VS, RL, LW, JI, HD, ZLL, LL, YSL, MWS, CMP, and WP performed the experiments. LH and HWL performed bacteria cultivation and verification as well as 16sRNA sample processing. BM and EFM analyzed 16sRNA data. TZ performed heart transplants.

## Abbreviations

Abx: antibiotics
APC: antigen presenting cell
Bifido: *Bifidobacterium pseudolongum*
CR: Cortical ridge
DC: Dendritic cell
Desulfo: *Desulfovibrio desulfuricans*
FRC: Fibroblastic reticular cell
HEV: High endothelial venule
IHC: Immunohistochemistry
ILC: Innate lymphoid cell
LN: Lymph node
MΦ: Macrophage
mLN: Mesenteric LN
pLN: Peripheral LN
SBT: Single bacterial strain transfer
Treg: Regulatory T cell

## ACKNOWLEDGEMENTS

This work was supported by National Institute of Health (NIH) National Heart, Lung and Blood Institute award R01HL148672 (JSB/BM), U01AI170050 (BM/JSB), and NIH National Institute of Allergy and Infectious Diseases training grant T32AI95190-10 (SJG). An earlier version of this manuscript has been released as a pre-print on bioRxiv https://doi.org/10.1101/2022.10.13.511915. EFM contributed to this work as an employee of the University of Maryland School of Medicine. The views expressed in this manuscript are his own and do not necessarily represent the views of the National Institutes of Health or the United States Government.

## DATA AVAILABILITY

The 16S rRNA gene sequencing data generated in this study was deposited to Genbank under BioProject PRJNA809764 (https://www.ncbi.nlm.nih.gov/bioproject/809764).

## COMPETING INTEREST STATEMENT

The authors have declared that no conflict of interest exists.

## Notes

### Competing Interest Statement

The authors have declared no competing interest.

### Summary of Updates

This updated manuscript has been streamlined in terms of the presentation of results and re-organized for clarity. Results also include additional experimental replicates for animals undergoing abx, SBT, transplant, and immunosuppression treatment for 40 days.

