## Supplemental Materials and Methods for "Early immunomodulatory program triggered by pro-tolerogenic *Bifidobacterium pseudolongum* drives cardiac transplant outcomes"

### Mice

Female C57BL/6 mice were used as cardiac transplant recipients, normal stool donors, and pregnant stool donors. C57BL/6 mice were purchased from The Jackson Laboratory (Bar Harbor, ME). Cardiac donors were female BALB/c mice, purchased from The Jackson Laboratory (Bar Harbor, ME). C57BL/6 and BALB/c mice were used between 8 and 14 weeks of age. TEa T cell receptor transgenic mice[54] on a C57BL/6 background that spontaneously develop colitis, due to a lack of Tregs, were used as the source for colitic stool and were a gift from Alexander Rudensky (Memorial Sloan Kettering Cancer Center, New York, NY, USA)[55,56]. All the procedures involving mice were performed in accordance with the guidelines and regulations set by the Office of Animal Welfare Assurance of the University of Maryland School of Medicine, under the approved IACUC protocol nos. 1518004 and 0121001.

### Stool specimen collection and DNA extraction

Stool pellets were collected fresh from cardiac transplant recipients at weekly or biweekly intervals until graft rejection or termination of the experiment. Stool samples were placed in RNAse-free 1.7-ml tubes (Denville Scientific, Holliston, MA) and then frozen at -80$^{o}$C until analyzed. DNA extraction from mouse stool pellets was adapted from procedures developed at the Institute for Genome Sciences and previously published[30]. In brief, 0.15-0.25 grams of fecal samples were extracted using the MagAttract PowerMicrobiome DNA/RNA kit (Qiagen, Hilden, Germany) implemented on a Hamilton STAR robotic platform and after a bead-beating step on a TissueLyzer II (Qiagen) in 96-deep well plates. Negative extraction controls were included to ensure that no exogenous DNA contaminated the samples during extraction. DNA quality control/quality assurance was performed using Nanodrop spectrophotometric measurements (ThermoFisher Scientific) and gel electrophoresis.

**16S rRNA gene amplicon sequencing**

Using a dual-indexing strategy for multiplexed sequencing developed at the Institute for Genome Sciences and described in detail elsewhere[57,58], the V3-V4 hypervariable region of the 16S rRNA gene was PCR amplified and sequenced on the Illumina MiSeq (Illumina, San Diego, CA). PCR reactions were set up using 319F (5′ - ACTCCTACGGGAGGCAGCAG - 3′) and 806R (5′ - GGACTACHVGGGTWTCTAAT - 3′) as universal primers, each with a linker sequence required for Illumina MiSeq 300bp paired-end sequencing, and a 12-bp heterogeneity-spacer index sequence to minimize biases associated with low-diversity amplicon sequencing[57,58]. Reactions were performed using Phusion High-Fidelity DNA polymerase (ThermoFisher Scientific). Successful amplification was confirmed using gel electrophoresis, after which amplicon cleanup and normalization was performed using the SequalPrep Normalization Plate kit (Invitrogen).

**Microbiota analysis**

After screening 16S rRNA gene reads for low-quality bases and short read lengths[57,58], paired-end read pairs were assembled using PANDAseq[59], demultiplexed, trimmed of artificial barcodes and primers, and assessed for chimeras using UCHIME in *de novo* mode implemented in Quantitative Insights Into Microbial Ecology (QIIME; release v. 1.9.1)[60]. Resulting quality trimmed sequences were then clustered *de novo* into OTUs with the GreenGenes 16S database (v. 13.8) in QIIME, with a minimum confidence threshold of 0.97. Taxonomic ranks were assigned to each sequence using the Ribosomal Database Project[61] Naïve Bayesian Classifier v.2.2[62] trained on the Greengenes database (Aug 2013 version)[63], using a confidence value cutoff of 0.8. To account for uneven sampling depth and to ensure less biases than the standard approach (total sum normalization), data were normalized with metagenomeSeq's cumulative sum scaling (CSS)[64] when appropriate. Sequencing analyses includes denoising, *de novo*, and referencebased chimera detection were conducted with UCHIME v5.1[65]. R packages of Vegan[66], Phyloseq[67], Bioconductor[68], metagenomeSeq[64] and ggplot2[69] were used for the following analyses. α diversity (within-sample diversity) was estimated using the Shannon diversity index[70]. β-Diversity (between-sample diversity) was determined through multidimensional scaling plots using Jensen–Shannon divergence distance to visualize the level of divergence at the different sampling times[71,72].

**Bacterial strain cultivation**

We previously described *B. pseudolongum* isolation methods from mouse feces[35], from which the *B. pseudolongum* UMB-MBP-01 (*Bifido* MD) strain was isolated. *B. pseudolongum subsp*. *pseudolongum Mitsuoka* ATCC25526 (*Bifido* ATCC) isolate was purchased from the American Type Culture Collection (ATCC, Manassas, VA). Both *B. pseudolongum* UMB-MBP-01 and ATCC25526 isolates were used in cell stimulation and cytokine assays, after being initially grown anaerobically at 37°C for 3–5 days on Bifidus Selective Media (BSM) agar plates (Millipore Sigma, Burlington, MA), from which a single colony was selected and grown in BSM broth (Millipore Sigma, 90273-500G-F) until stationary phase (up to 3 days). *Desulfovibrio desulfuricans subsp. desulfuricans* (ATCC 27774) was purchased from ATCC (Manassas, VA) and grown in ATCC 1249 Modified Baar's Medium (MBM) for sulfate reducers, which was made according to the ATCC protocol. Cultures were initially incubated under anaerobic conditions for 5 days on MBM agar plates, after which single colonies were chosen, transferred to liquid media, and incubated for up to 3 weeks.

**Fecal microbiota transplant (FMT) and Single Strain Bacteria Transfer (SBT)**

As previously reported[30], mice were first fed abx (kanamycin, gentamicin, colistin, metronidazole, and vancomycin) ad libitum in drinking water on days –6 through –1, and then

FMT or SBT was performed on day 0 by oral gavage. Cultured *B. pseudolongum* ATCC25526, *B. pseudolongum* UMB-MBP-01, and *D. desulfuricans* ATCC 27774 bacteria were administered in 100 µl (1 at $10^8$ bacteria/ml) in PBS and used fresh. Fecal samples were administered in 200 µl (200 □g/ml in PBS). On day 0, C57BL/6 mice also received BALB/c heart transplants. Mice received daily immunosuppression with tacrolimus (3 mg/kg/d s.c.) starting on day 0[73,74].

**Reagents**

Antibiotics were USP grade or pharmaceutical secondary standard: kanamycin sulfate (0.4 mg/ml, MilliporeSigma), gentamicin sulfate (0.035 mg/ml, MilliporeSigma), colistin sulfate (850 U/ml, MilliporeSigma), metronidazole (0.215 mg/ml, MilliporeSigma), and vancomycin hydrochloride (0.045 mg/ml, MilliporeSigma) were dissolved in vivarium drinking water and administered ad libitum. Tacrolimus (USP grade, MilliporeSigma) was reconstituted in DMSO (USP grade, MilliporeSigma) at 20 mg/ml. DMSO stock was diluted to 2–3 mg/kg with absolute ethanol (USP grade, Decon Labs). DMSO/ethanol stock was diluted 1:5 in sterile PBS for s.c. injection and injected at 10 µl/g.

**Cell culture**

Murine bone marrow (BM) derived dendritic cells (BMDC) were generated as previously described[75]. Briefly, bone marrow (BM) cells of wild-type mice were treated with 10 ng/mL GM-CSF (R&D Systems, Minneapolis, MN) for 10 days in Petri dishes, and the loosely attached cells were collected and CD11c+ DC were purified by CD11c-positive selection kit (Stemcell Technologies). Peritoneal MF were harvested as previously described[76]. Briefly, MΦ were collected 4 days after i.p. injection of Remel Thioglycollate solution (Thermo Fisher Scientific, Waltham, MA). Cells were resuspended in complete medium and stimulated as described below.

**Cell stimulation and cytokine assays**

*D. desulfuricans* ATCC 27774 or *B. pseudolongum* cultures were spun down, the supernatant collected, and bacterial cells killed by UV exposure at 100μJ/cm$^2$ for four 15-minute cycles with a UV CrossLinker (Fisher Scientific, Hampton, NH). BMDC and peritoneal MF (1 x 10$^6$) were seeded into duplicate 12-well plates in 1 ml complete medium. The following day, myeloid cells were stimulated with media alone or UV-killed *B. pseudolongum* MD or *D. desulfuricans* bacteria at a multiplicity of infection (MOI) of 5 for 24 hours. Culture supernatants were collected for ELISA. ELISAs for TNFα, IL-6, and IL-10 were purchased from BioLegend (San Diego, CA) and CCL19 from Boster Biological Technology (Pleasanton, CA).

**Histology**

Cardiac allografts were assessed for survival by abdominal palpation; harvested at day 40, day 60, or time of rejection defined by two consecutive days without palpable heart beats; fixed for 2 days in 10% buffered formalin (ThermoFisher Scientific, Waltham, MA); and transferred to 70% ethanol. Samples were embedded in paraffin, processed, and stained with H&E or Masson's trichrome. H&E samples were evaluated for parenchymal rejection score (from 0–4) using previously published criteria[77]. Trichrome samples were scored 1–4, with 1 as minimal (<10% area), 2 as mild (10%–30%), 3 as moderate (30%–50%), and 4 as severe (>50%). Slides were read by multiple individuals blinded to the treatment group of the specimen. Scores for each slide were averaged. Inflammation scores were calculated by combining the averaged trichrome and H&E score from each sample.

**Immunohistochemistry**

Peripheral LN, mesenteric LN, and small bowel segments (duodenal-jejunal junction) were excised and washed in cold PBS before freezing in OCT (Sakura Finetek, Torrance, CA) and then stored at -80°C. LN cryosections were cut in triplicate at 6 μm for LN and 10 μm for intestine using a Microm HM 550 cryostat (Thermo Fisher Scientific, Waltham, MA). Sections attached to slides were fixed with cold acetone/ethanol (1:1) solution (Lonza, Morristown, NJ) and then blocked with 10% donkey and/or goat serum. Primary and secondary antibodies are listed in **Supplementary Table 1.** Slides were then fixed with 4% paraformaldehyde/PBS (Alfa Aesar, Haverhill, MA) and mounted with Prolong Gold Antifade Mountant with or without DAPI (Thermo Fisher Scientific, Waltham, MA). Images were acquired using an Accu-Scope EXC-500 fluorescent microscope (Nikon, Melville, NY) and analyzed with Volocity image analysis software (PerkinElmer, Waltham, MA). Mean of mean fluorescence intensity (MFI) was calculated within demarcated high endothelial venules (HEV) and cortical ridge (CR) regions of mLN and pLN as well as of whole intestinal images. Percent area was calculated by dividing the sum area of demarcated regions with marker fluorescence greater than a given threshold divided by total area analyzed. Treatment groups were compared using quantitation of MFI multiplied by percent area to express both area and intensity of cell and ECM markers. MFI and percent area were quantified based on at least 2 independent experiments with 3 mice/group, 3 mLN or pLN/mouse or 2 intestine segments/mouse, 3 sections/LN or segment of intestine (duodenal-jejunal junction), and 3–5 fields imaged/section, i.e., 20-30 total microscopy fields per condition over 2 experiments or 30-45 total microscopy fields per condition over 3 experiments.

**Flow cytometry**

Cells were passed through 70-μm nylon mesh screens (Thermo Fisher Scientific, Waltham, MA) to produce single-cell suspensions. Cell suspensions were treated with anti-CD16/32 (clone 93, eBioscience) to block Fc receptors, and then stained for 30 minutes at 4°C with antibodies against surface molecules (**Supplementary Table 1**) and washed 2 times in FACS buffer [phosphate buffered saline (PBS) with 0.5% w/v Bovine serum albumin (BSA).

Samples were analyzed with an LSR Fortessa Cell Analyzer (BD Biosciences), and data analyzed with FlowJo software version 10.6 (BD Biosciences).

**Heat Map Construction (Immunohistochemistry and flow cytometry data)**

Qualitative heat maps were designed to visually denote statistically significant increases or decreases in IHC and flow cytometry markers for a given treatment relative to control. Maps were designed using Prism Graphpad software. Increased marker expression was entered as "+1" (red box) and decreased expression was entered as "-1" (blue box) while no change was entered as "0" (white box).

**Statistics**

Datasets were analyzed using GraphPad Prism 9.3.1 (San Diego, CA) with statistical significance defined for $P < 0.05$. For comparisons of fluorescent markers (including laminin $\alpha 4$:$\alpha 5$ ratios), serum markers, and inflammation scores, Turkey's multiple comparisons tests of one-way ANOVA were used to test for significance.
