## Supplemental Figures for "Early immunomodulatory program triggered by pro-tolerogenic *Bifidobacterium pseudolongum* drives cardiac transplant outcomes"

**Supplemental Figure S1. Distribution of the most abundant taxonomic groups in gut microbiota.** Unclassified, abundant bacterial taxonomic groups at the species level merged at the genus level for easy visualization.

**Supplemental Figure S2.** Mice given abx (6 days), then SBT (*PBS, Bifido MD,* or *Desulfo*). After 3 days, LN, spleen, and intestinal segments harvested. **(a)** Flow cytometry of mLN CD11c+ DCs and F4/80+ MΦ; pLN B220+ B cells and CD11c+ DC*.* IHC for **(b)** mLN Foxp3+ Tregs, CD11c+ DC, and F4/80+ MΦ; **(c)** pLN Foxp3+ Tregs, CD11c+ DC, and F4/80+ MΦ*;* **(d)** laminin $\alpha$4 and $\alpha$5 and ratios in mLN and pLN; and **(e)** intestine Foxp3+ Tregs, CD11c+ DC, and F4/80+ MΦ. Ordinary one-way ANOVA with Tukey's multiple comparisons test. * $p < 0.05$; ** $p < 0.01$, *** $p < 0.001$, **** $p < 0.0001$.

**Supplemental Figure S3.** Mice treated with abx (6 days), then SBT (*PBS, Bifido MD,* or *Desulfo*), followed by tacrolimus for 2 days. After 3 days, LN and intestinal segments harvested for flow and IHC. Flow cytometry of **(a)** mLN B220+ B cells, CD4+ T cells, CD8+ T cells, Foxp3+ Tregs, F4/80+ MΦ, and CD11c+ DC; and **(b)** pLNs analyzed for B220+ B cells, CD4+ T cells, CD8+ T cells, Foxp3+ Tregs, F4/80+ MΦ, and CD11c+ DC. IHC for **(c)** mLN CR Foxp3+ Tregs, CD11c+ DC, and F4/80+ MΦ; **(d)** pLN Foxp3+ Tregs, CD11c+ DC, F4/80+ MΦ*,* and pLN laminin $\alpha$4 and $\alpha$5 ratios; and **(e)** intestine Foxp3+ Tregs, CD11c+ DC, and F4/80+ MΦ. 3 mice/group. At least 2 mLN and pLN/mouse. Ordinary one-way ANOVA with Tukey's multiple comparisons test. * $p < 0.05$; ** $p < 0.01$, *** $p < 0.001$, **** $p < 0.0001$.

**Supplemental Figure S4.** Mice given abx (6 days), then SBT (PBS, *Bifido* MD, or *Desulfo*) on the day of heart transplant with injection of TEa T cells (2 x $10^5$ cells i.v.), followed by tacrolimus for 7 days. IHC for **(a)** laminin $\alpha 4$ and $\alpha 5$ and ratios in mLN as well as intestine Foxp3+ Tregs and CD11c+ DC. **(b-e)** Mice given abx (6 days), then SBT (PBS, *Bifido* MD, or *Desulfo*) on the day of heart transplant with injection of TEa T cells (2 x 105 cells i.v.), followed by tacrolimus for 14 days. IHC for **(b)** mLN Foxp3+ Tregs, CD11c+ DC, and F4/80+ MΦ*;* **(c)** pLN Foxp3+ Tregs, CD11c+ DC, and F4/80+ MΦ*;* **(d)** laminin $\alpha 4$ and $\alpha 5$ and ratios in mLN and pLN; and **(e)** intestine Foxp3+ Tregs, CD11c+ DC, and F4/80+ MΦ*.* Ordinary one-way ANOVA with Tukey's multiple comparisons test. * $p < 0.05$; ** $p < 0.01$, *** $p < 0.001$, **** $p < 0.0001$.

**Supplemental Figure S5. Effect of *Bifido* and *Desulfo* on cardiac transplant inflammation and LN remodeling.** Mice given abx (6 days), then SBT (PBS, *Bifido* MD, or *Desulfo*) on the day of heart transplant with injection of TEa T cells (2 x $10^5$ cells i.v.), followed by tacrolimus for 7 or 14 days. **(a)** Cardiac allograft H&E at 7 and **(b)** 14 days. **(c)** Representative H&E images at 7 days, 20x. 3 mice/group, 3 sections/graft, 2 fields/section. **(d)** Serum analyzed following treatment with abx (6 days), then SBT, followed by tacrolimus (3 mg/kg/d s.c.) for 7 days using clinical tandem mass spectrometry. Three mice per *Bifido* and Desulfo treatment groups, 4 mice treated with normal SBT. Ordinary one-way ANOVA with Tukey's multiple comparisons test. * $p < 0.05$; ** $p < 0.01$, *** $p < 0.001$, **** $p < 0.0001$.

**Supplemental Figure S6.** Mice given abx (6 days), then SBT (WT stool, *Bifido ATCC,* or *Desulfo*) on the day of heart transplant, followed by tacrolimus for 40 days. IHC for **(a)** Foxp3+ Tregs, CD11c+ DC; and F4/80+ MΦ around HEV and in CR of mLN; **(b)** Foxp3+ Tregs, CD11c+ DC, and F4/80+ MΦ around HEV and in CR of pLN; and **(c)** laminin $\alpha 4$ and $\alpha 5$ ratios in pLN and mLN. Three mice per group, at least 2 mLN, pLN, and sections of intestine at duodenal-jejunal junction per mouse, 3 sections/LN/staining panel. Ordinary one-way ANOVA with Tukey's multiple comparisons test. * $p < 0.05$; ** $p < 0.01$, *** $p < 0.001$, **** $p < 0.0001$.

**Supplemental Figure S7.** Mice given abx (6 days), SBT (WT stool, *Bifido MD,* or *Desulfo*) on the day of heart transplant, followed by tacrolimus for 60 days. Flow cytometry for **(a)** CD4+ T cells and CD8+ T cells in spleen; and pLN F4/80+ M$\Phi$. IHC of **(b)** Foxp3+ Tregs, F4/80+ M$\Phi$, and CD11c+ DC in mLN; **(c)** Foxp3+ Tregs, F4/80+ M$\Phi$, and CD11c+ DC in pLN; **(d)** laminin $\alpha$4:$\alpha$5 ratios in mLN and pLN; and **(e)** Foxp3+ Tregs, CD11c+ DC, and F4/80+ M$\Phi$ in intestine. Ordinary one-way ANOVA with Tukey's multiple comparisons test. * $p < 0.05$; ** $p < 0.01$, *** $p < 0.001$, **** $p < 0.0001$.

**Supplemental Table 1: List of Antibodies**

| Specificity | Clone | Catalog No. |
|---|---|---|
| Donkey anti-mouse IgG-AF647 | Polyclonal | Jackson ImmunoResearch; 715-605-151 |
| Goat anti-rabbit AF488 | Polyclonal | Jackson ImmunoResearch; 111-545-003 |
| Goat anti-rabbit Cy5 | Polyclonal | Jackson ImmunoResearch; 111-175-003 |
| Donkey anti-rabbit DL405 | Polyclonal | Jackson ImmunoResearch; 711-476-152 |
| Donkey anti-rabbit IgG-AF488 | Polyclonal | Jackson ImmunoResearch; 711-545-152 |
| Goat anti-rabbit IgG-AF594 | Polyclonal | Jackson ImmunoResearch; 111-585-003 |
| Donkey anti-rabbit IgG-AF647 | Polyclonal | Jackson ImmunoResearch; 711-606-152 |
| Donkey anti-rat CY3 | Polyclonal | Jackson ImmunoResearch; 712-166-153 |
| Goat anti-rat Cy5 | Polyclonal | Jackson ImmunoResearch; 112-175-167 |
| Goat anti-rat IgG AF594 | Polyclonal | Jackson ImmunoResearch; 112-586-143 |
| Rat anti-B220 | RA3-6B2 | eBioSc; 17-0452-82 |
| Armenian hamster anti-CD103 | 2E7 | Biolegend; 121422 |
| Rat anti-CD11b | M1/70 | eBioSc; 11-0112-81 |

| | | |
|---|---|---|
| Hamster anti-mouse CD11c | HL3 | BD; 550283 |
| Hamster anti-mouse CD11c FITC | HL3 | BD; 553801 |
| Rat anti-CD163 | S150491 | Biolegend; 155307 |
| Rat anti-CD206 | C068C2 | Biolegend; 141717 |
| Rat anti-CD4 | GK1.5 | Biolegend; 100401 |
| Rat anti- mouse CD40 | 3/23 | Biolegend; 124609 |
| Rat anti-CD44 | IM7 | eBioSc; 17-0441-82 |
| Hamster anti-mouse CD69 | H1.2F3 | BD; 553237 |
| Rat anti-CD8 | 53-6.7 | Biolegend; 100701 |
| Armenian hamster anti-mouse CD80 | 16-10A1 | Biolegend; 104705 |
| Rat anti-mouse CD86 | GL-1 | Biolegend; 105011 |
| Rat anti-ER-TR7 | ER-TR7 | Santa Cruz; SC-73355 |
| Rat anti-ER-TR7 | ER-TR7 | Novus; NB100-64932AF405 |
| Rat anti-F4/80 | BM8 | eBioSc; 11-4801-81 |
| Rat anti-Foxp3 | PCH101 | eBioSc; 14-4776-82 |
| Rat anti-mouse Laminin a4 | 775830 | R&D; MAB3837 |
| Rabbit anti-mouse Laminin a5 | Polyclonal | Novus Biol; NBP1-18714 |
| Rat anti-mouse MHC II (I-A/I-E) | M5/114.15.2 | Biolegend; 107619 |

### Suppl. Fig. S1

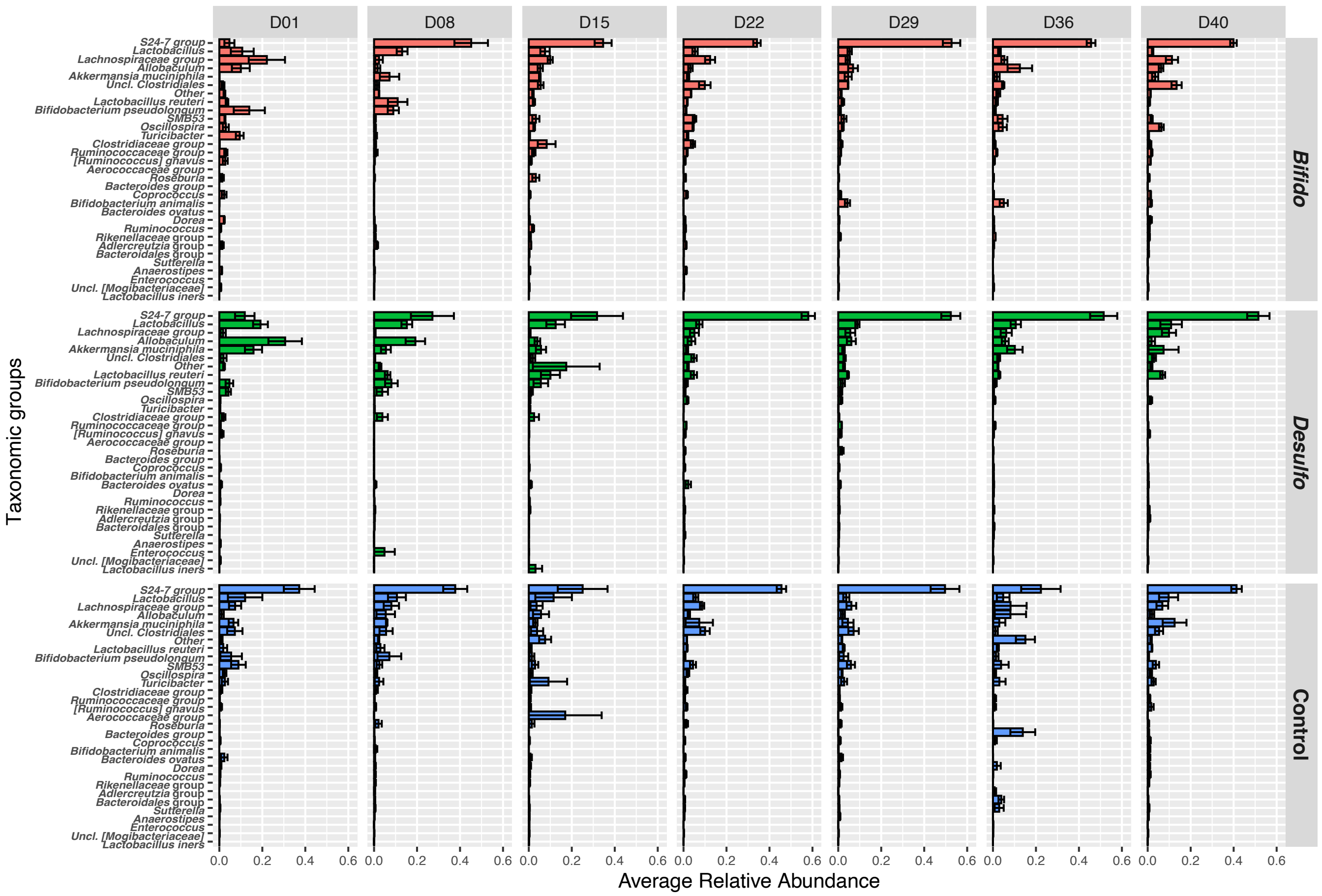

D01
D08
D15
D22
D29
D36
D40
Bifido
Desulfo
Control
Taxonomic groups
Average Relative Abundance
S24-7 group
Lactobacillus
Lachnospiraceae group
Allobaculum
Akkermansia muciniphila
Uncl. Clostridiales
Other
Lactobacillus reuteri
Bifidobacterium pseudolongum
SMB53
Oscillospira
Turicibacter
Clostridiaceae group
Ruminococcaceae group
[Ruminococcus] gnavus
Aerococcaceae group
Roseburia
Bacteroides group
Coprococcus
Bifidobacterium animalis
Bacteroides ovatus
Dorea
Ruminococcus
Rikenellaceae group
Adlercreutzia group
Bacteroidales group
Sutterella
Anaerostipes
Enterococcus
Uncl. [Mogibacteriaceae]
Lactobacillus iners
0.0
0.2
0.4
0.6

### Suppl. Fig. S2

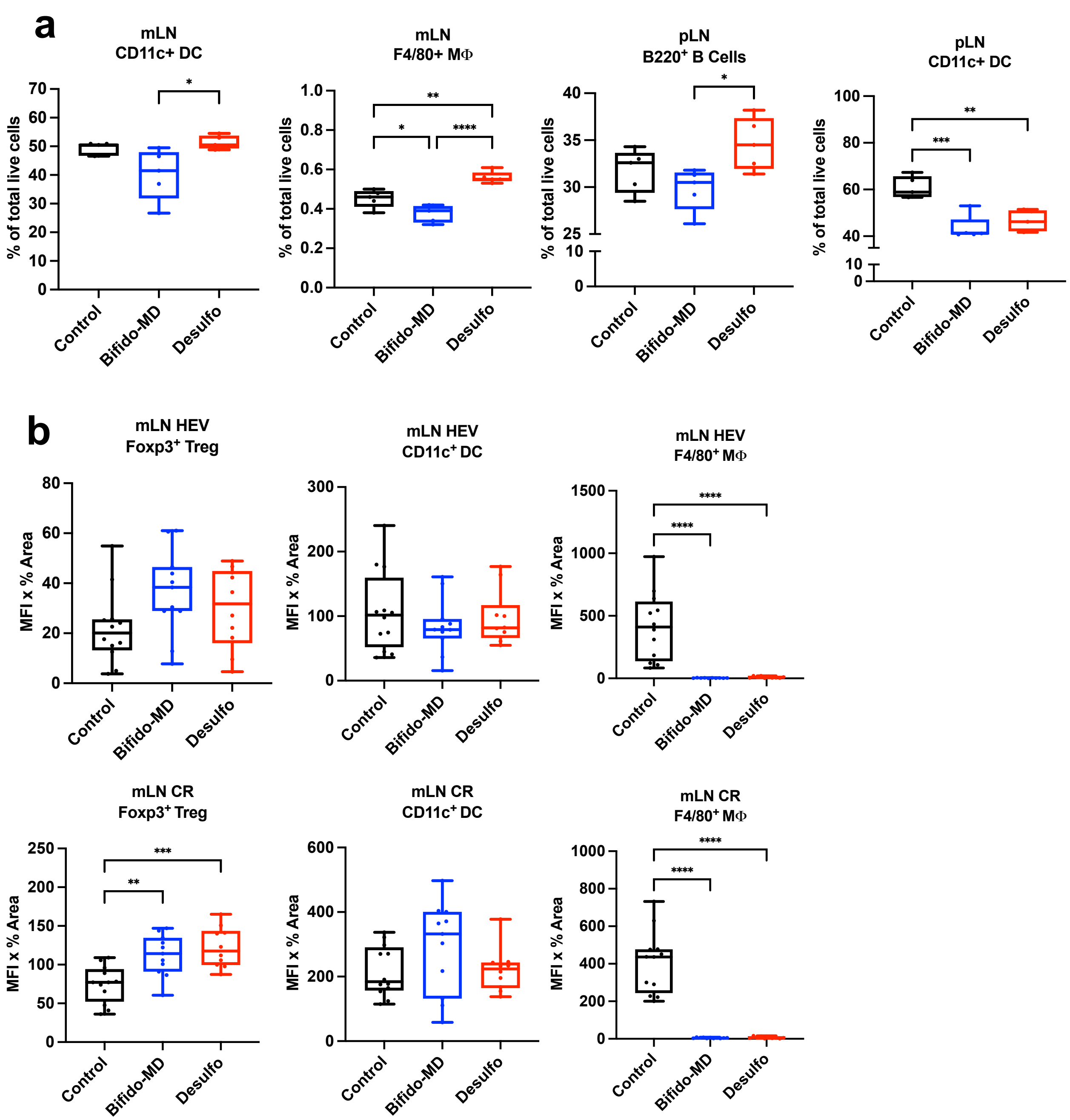
a
mLN CD11c+ DC
% of total live cells
mLN F4/80+ MΦ
pLN B220+ B Cells
pLN CD11c+ DC
Control
Bifido-MD
Desulfo
b
mLN HEV Foxp3+ Treg
mLN HEV CD11c+ DC
mLN HEV F4/80+ MΦ
mLN CR Foxp3+ Treg
mLN CR CD11c+ DC
mLN CR F4/80+ MΦ
MFI x % Area

### Suppl. Fig. S2

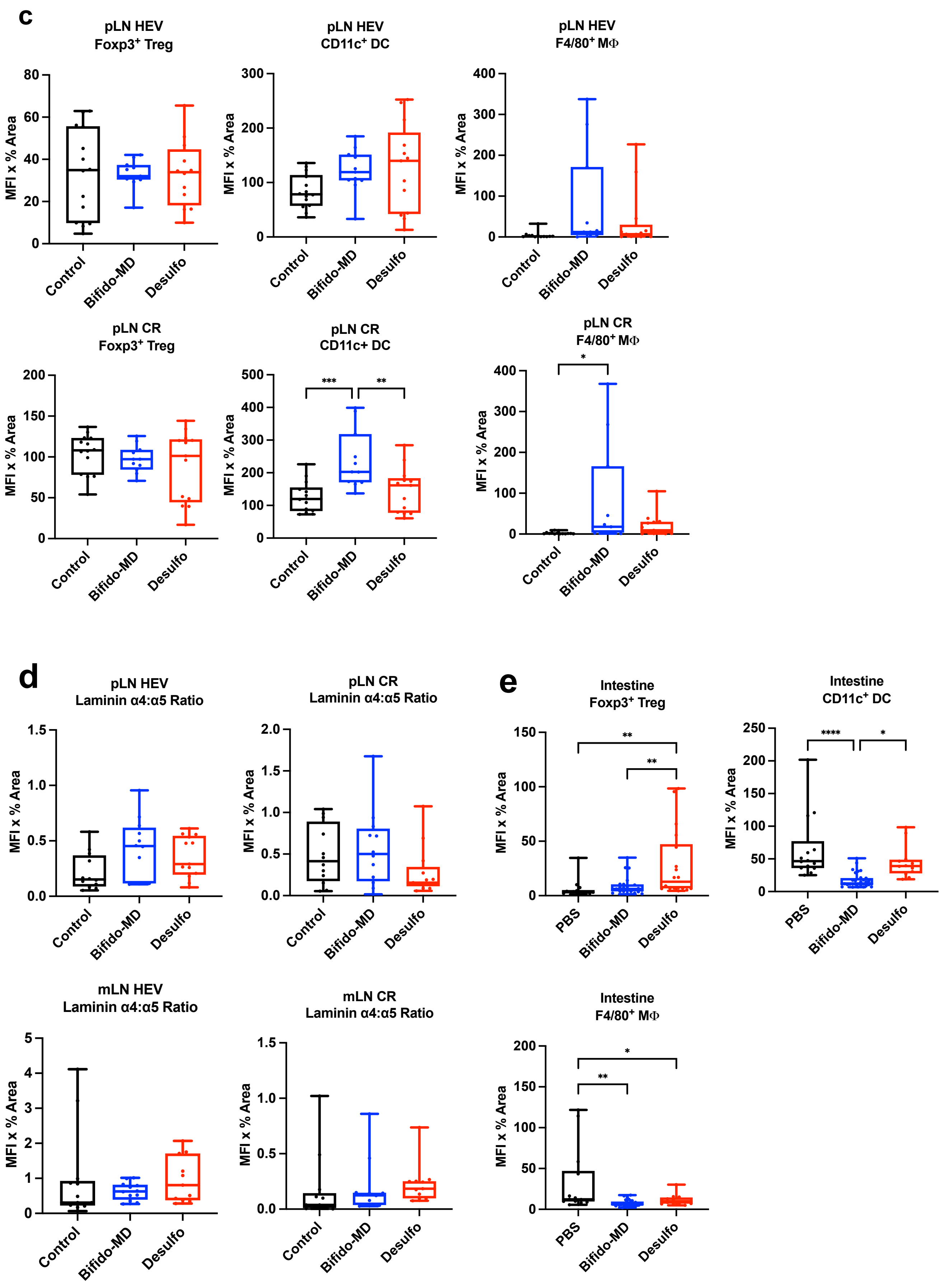

c
pLN HEV
Foxp3+ Treg
MFI x % Area
Control
Bifido-MD
Desulfo
pLN HEV
CD11c+ DC
pLN HEV
F4/80+ MΦ
pLN CR
Foxp3+ Treg
pLN CR
CD11c+ DC
***
**
pLN CR
F4/80+ MΦ
*
d
pLN HEV
Laminin α4:α5 Ratio
pLN CR
Laminin α4:α5 Ratio
mLN HEV
Laminin α4:α5 Ratio
mLN CR
Laminin α4:α5 Ratio
e
Intestine
Foxp3+ Treg
PBS
**
Intestine
CD11c+ DC
****
*
Intestine
F4/80+ MΦ

### Suppl. Fig. S3

#### Mesenteric LN

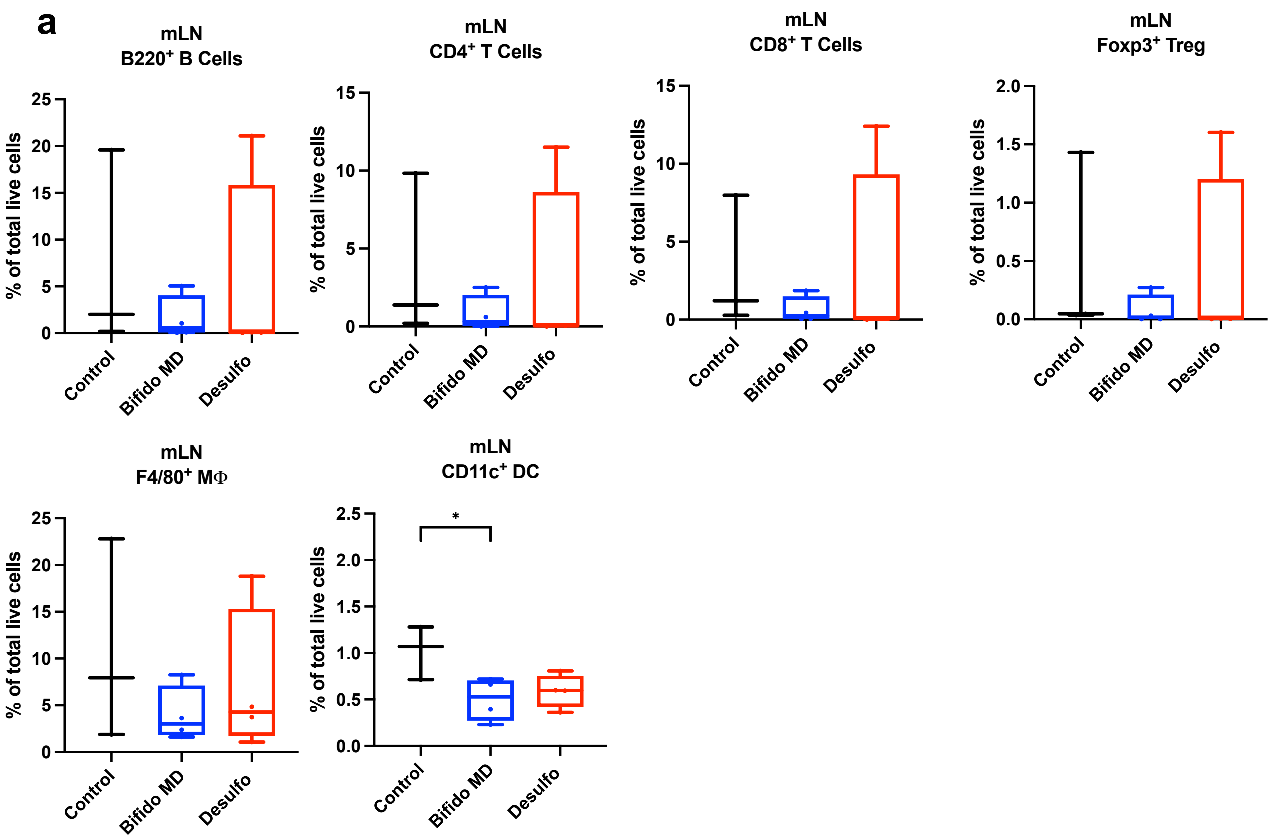


#### Peripheral LN

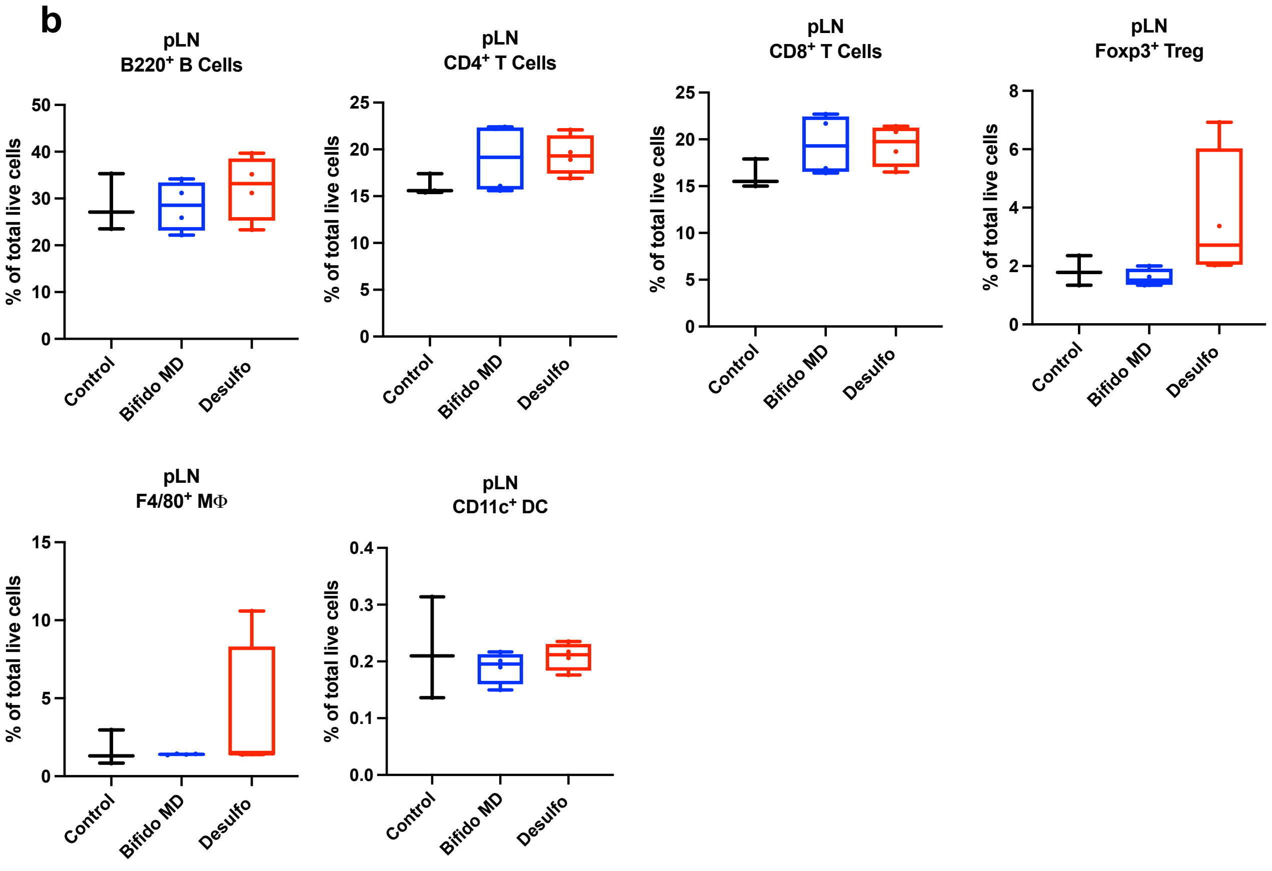

### Suppl. Fig. S3

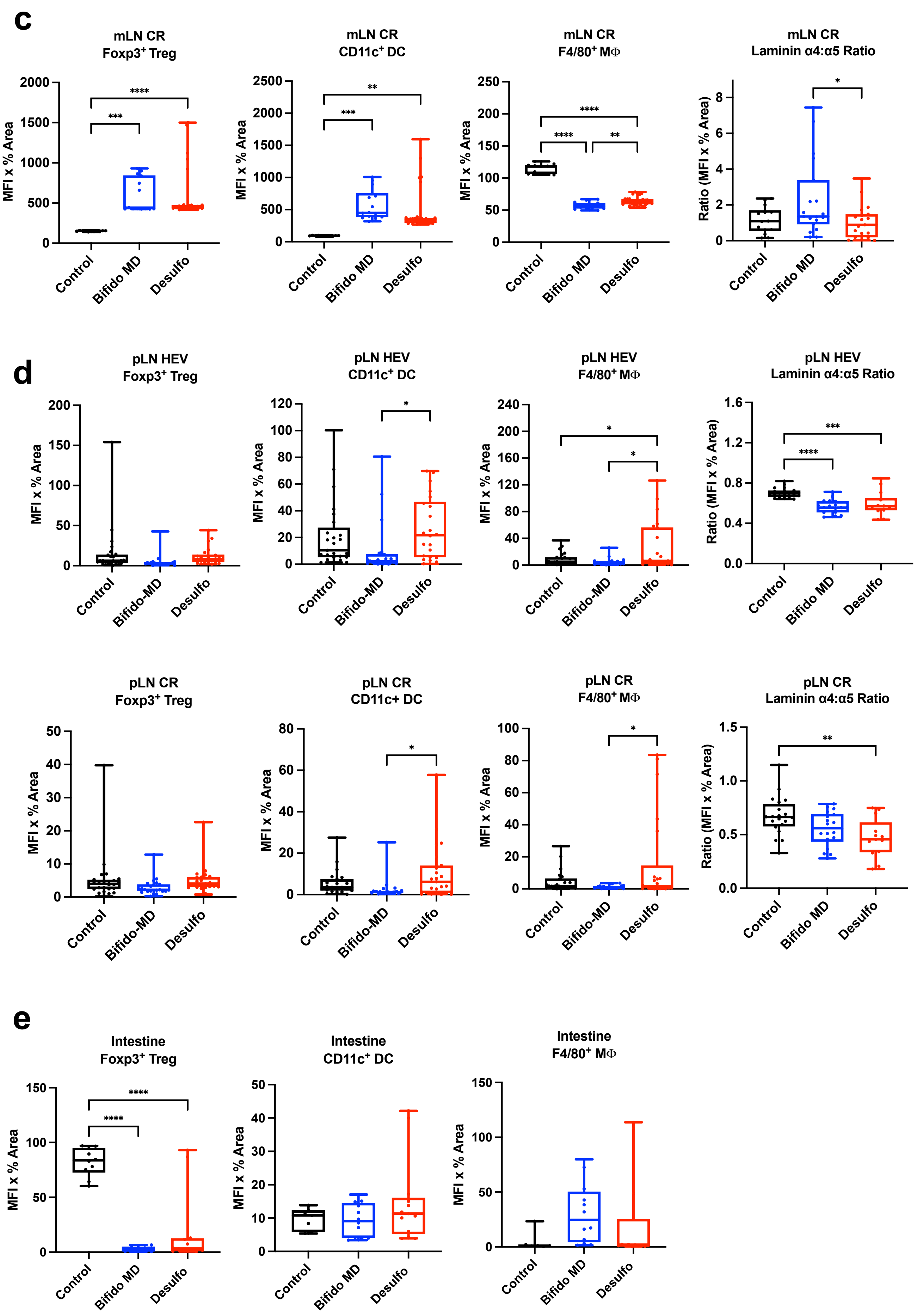
c
mLN CR
Foxp3+ Treg
MFI x % Area
Control
Bifido MD
Desulfo
mLN CR
CD11c+ DC
mLN CR
F4/80+ MΦ
mLN CR
Laminin α4:α5 Ratio
Ratio (MFI x % Area)
d
pLN HEV
Foxp3+ Treg
Bifido-MD
pLN HEV
CD11c+ DC
pLN HEV
F4/80+ MΦ
pLN HEV
Laminin α4:α5 Ratio
pLN CR
Foxp3+ Treg
pLN CR
CD11c+ DC
pLN CR
F4/80+ MΦ
pLN CR
Laminin α4:α5 Ratio
e
Intestine
Foxp3+ Treg
Intestine
CD11c+ DC
Intestine
F4/80+ MΦ

### Suppl. Fig. S4

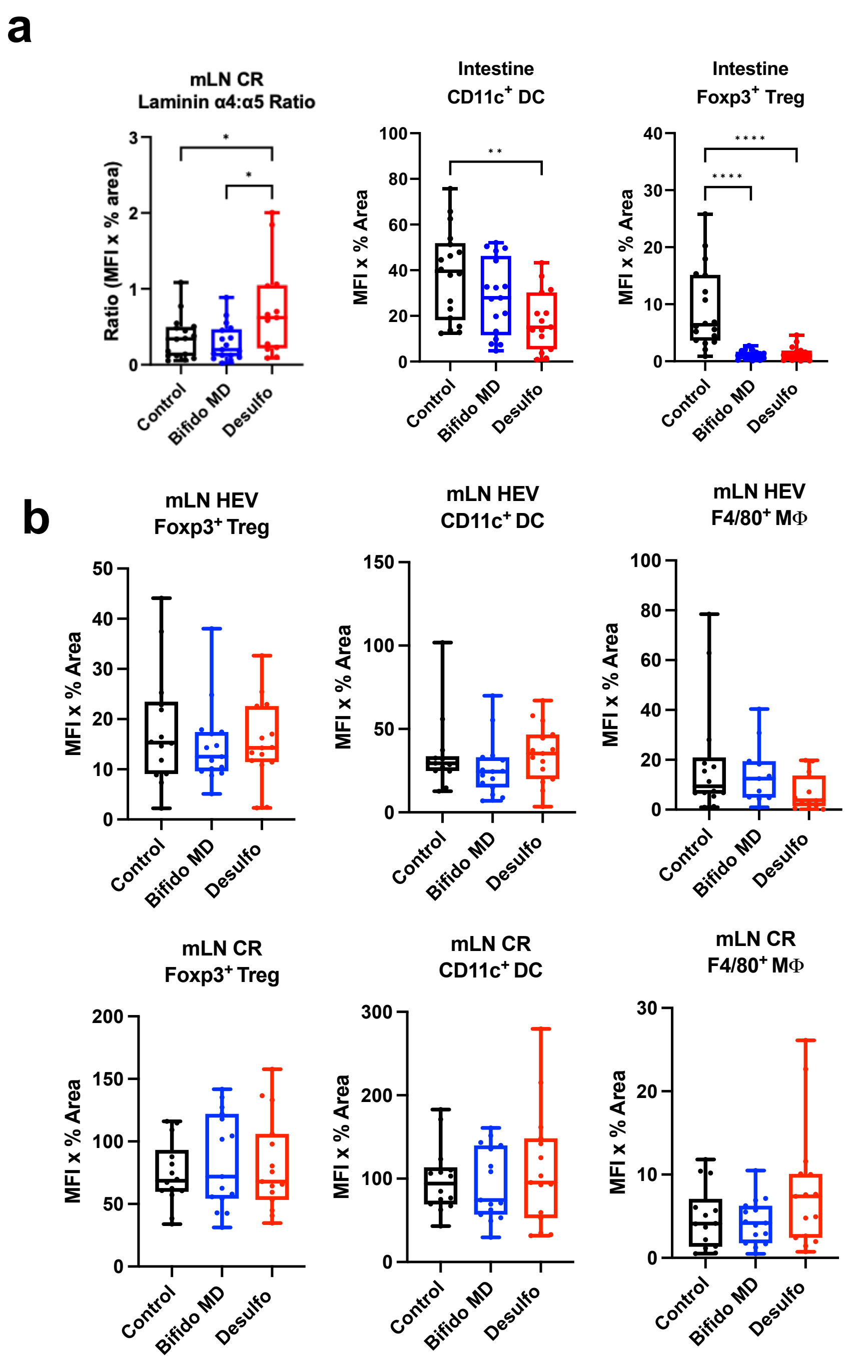
a
mLN CR
Laminin α4:α5 Ratio
Ratio (MFI x % area)
Intestine
CD11c+ DC
MFI x % Area
Intestine
Foxp3+ Treg
MFI x % Area
Control
Bifido MD
Desulfo
*
**
****
b
mLN HEV
Foxp3+ Treg
mLN HEV
CD11c+ DC
mLN HEV
F4/80+ MΦ
mLN CR
Foxp3+ Treg
mLN CR
CD11c+ DC
mLN CR
F4/80+ MΦ
MFI x % Area
Control
Bifido MD
Desulfo

### Suppl. Fig. S4

c

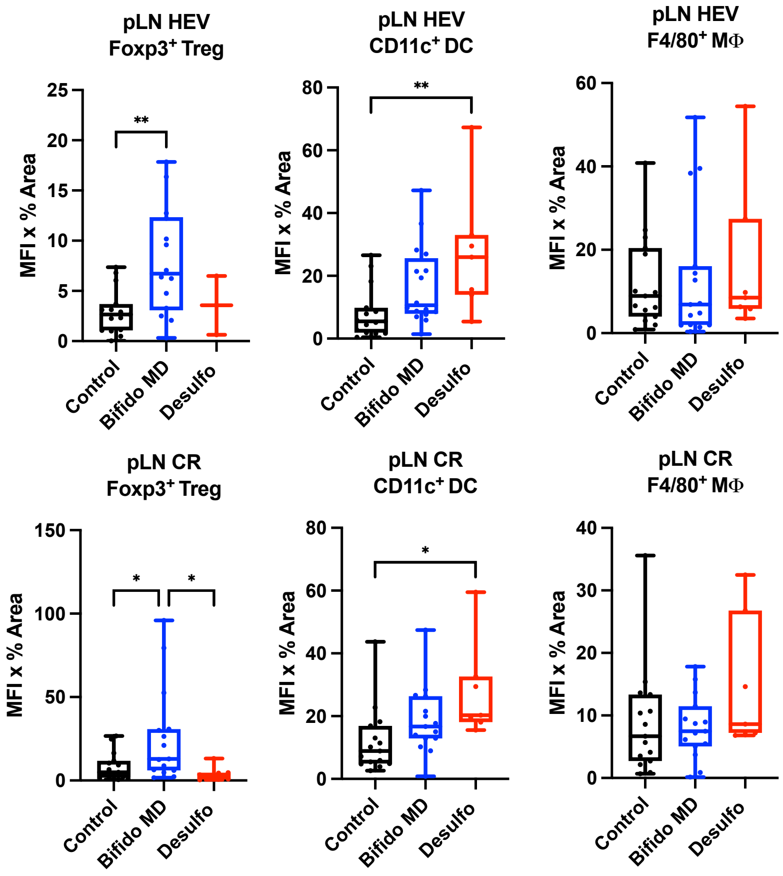


d

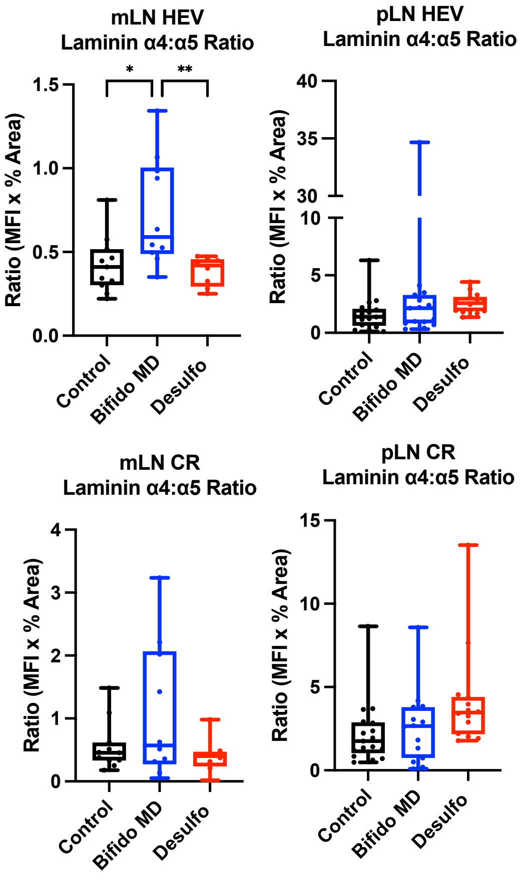


e

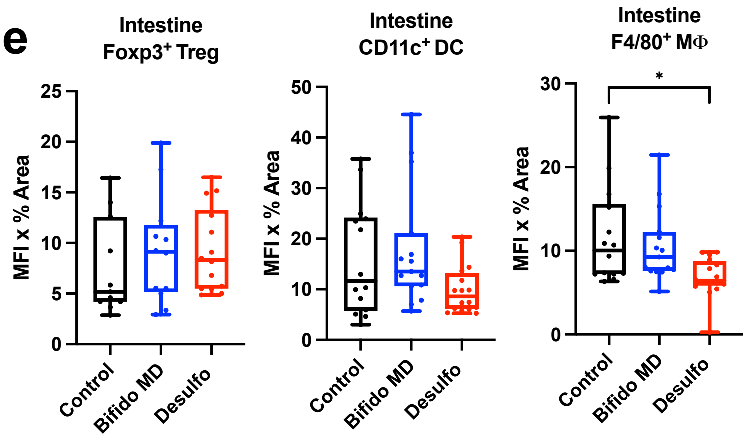

Suppl. Fig. S5

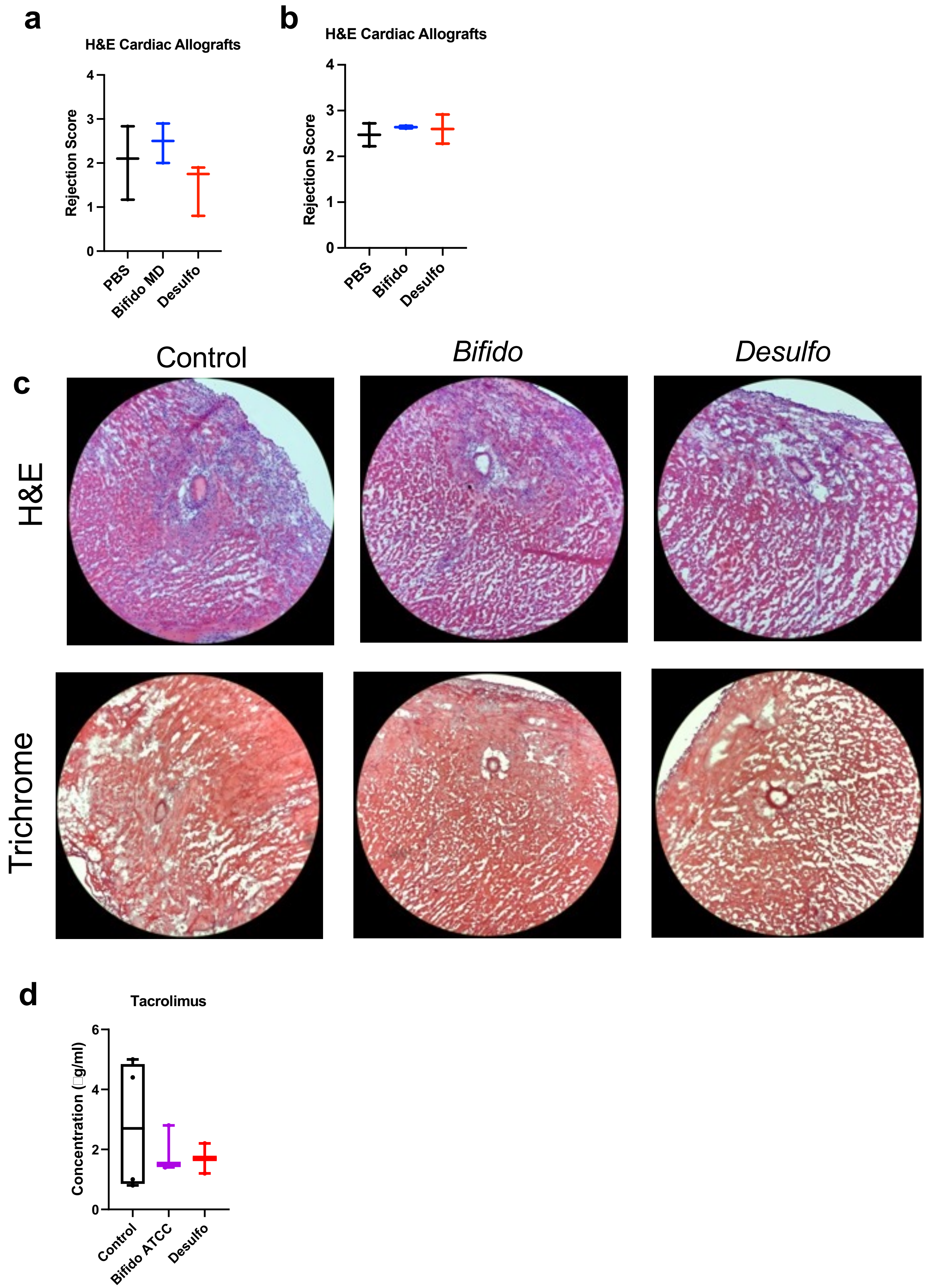

a
H&E Cardiac Allografts
Rejection Score
0
1
2
3
4
PBS
Bifido MD
Desulfo
b
H&E Cardiac Allografts
Rejection Score
0
1
2
3
4
PBS
Bifido
Desulfo
c
Control
Bifido
Desulfo
H&E
Trichrome
d
Tacrolimus
Concentration (g/ml)
0
2
4
6
Control
Bifido ATCC
Desulfo

### Suppl. Fig. S6

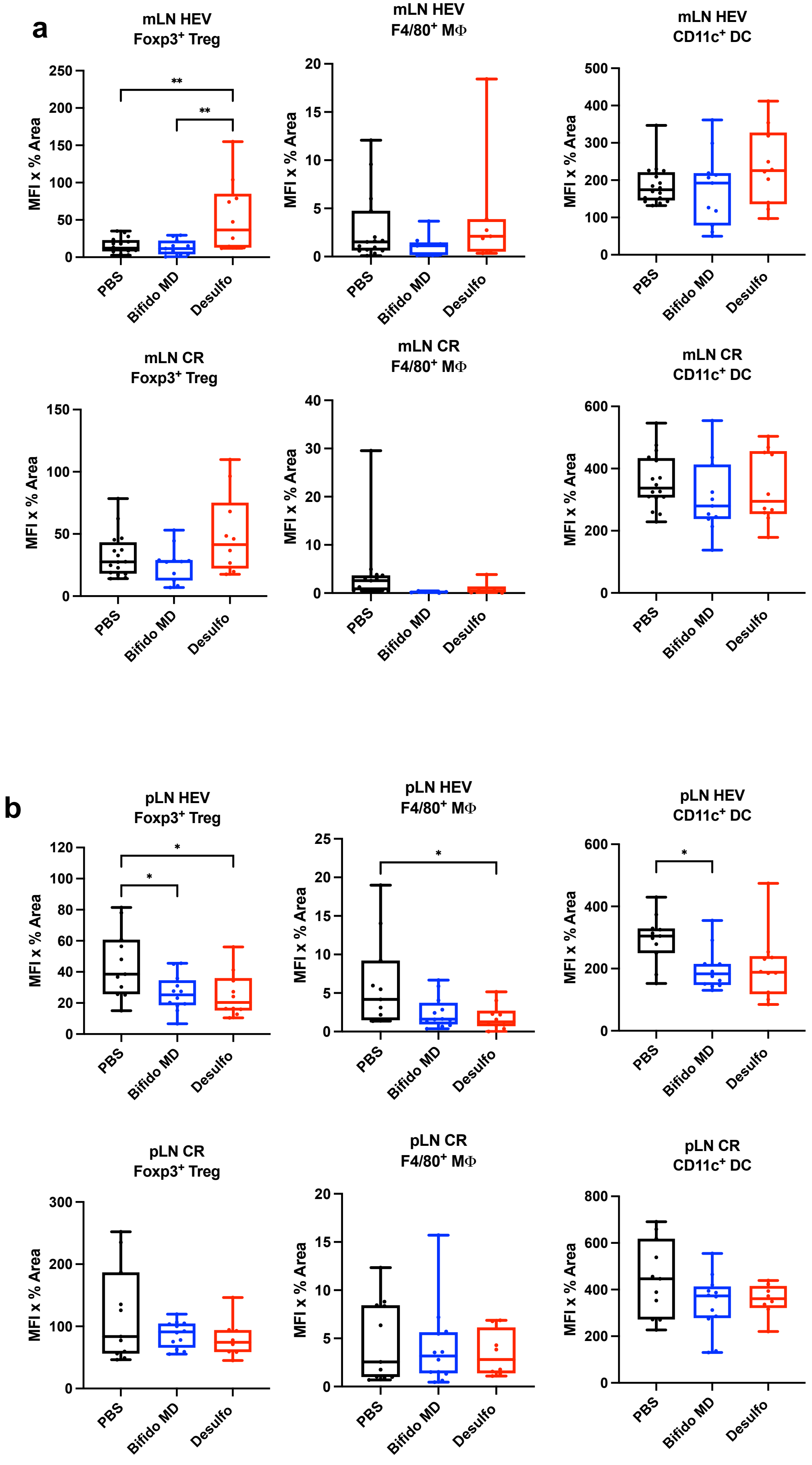

a
mLN HEV
Foxp3+ Treg
MFI x % Area
250
200
150
100
50
0
**
**
PBS
Bifido MD
Desulfo
mLN HEV
F4/80+ MΦ
20
15
10
5
0
mLN HEV
CD11c+ DC
500
400
300
200
100
0
mLN CR
Foxp3+ Treg
150
100
50
0
mLN CR
F4/80+ MΦ
40
30
20
10
0
mLN CR
CD11c+ DC
600
400
200
0
b
pLN HEV
Foxp3+ Treg
120
100
80
60
40
20
0
*
*
pLN HEV
F4/80+ MΦ
25
20
15
10
5
0
*
pLN HEV
CD11c+ DC
600
400
200
0
*
pLN CR
Foxp3+ Treg
300
200
100
0
pLN CR
F4/80+ MΦ
20
15
10
5
0
pLN CR
CD11c+ DC
800
600
400
200
0

### Suppl. Fig. S6

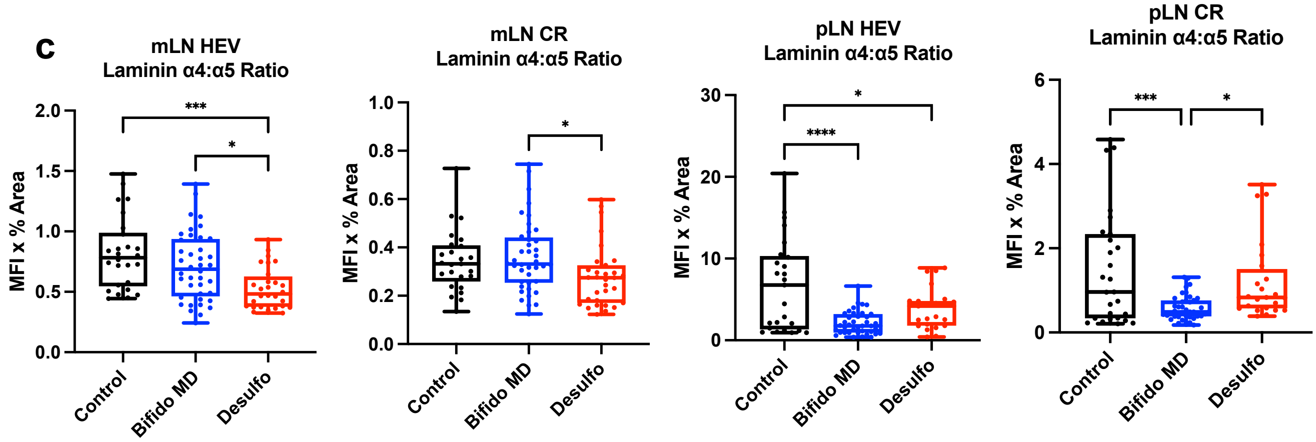

c
mLN HEV
Laminin α4:α5 Ratio
MFI x % Area
2.0
1.5
1.0
0.5
0.0
***
*
Control
Bifido MD
Desulfo
mLN CR
Laminin α4:α5 Ratio
1.0
0.8
0.6
0.4
0.2
0.0
pLN HEV
Laminin α4:α5 Ratio
30
20
10
0
****
pLN CR
Laminin α4:α5 Ratio
6
4
2

### Suppl. Fig. S7

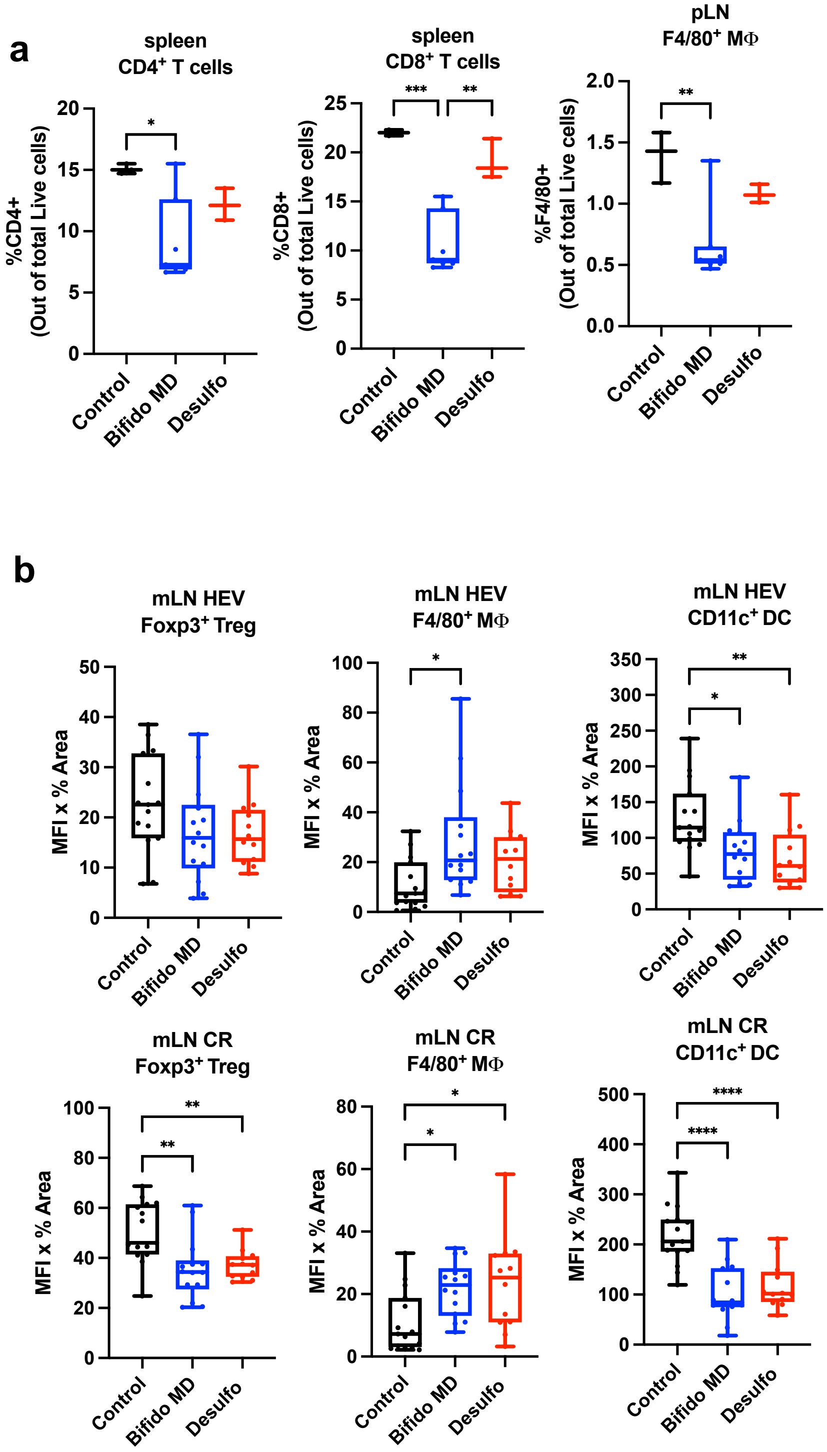

a
spleen
CD4+ T cells
%CD4+ (Out of total Live cells)
spleen
CD8+ T cells
%CD8+ (Out of total Live cells)
pLN
F4/80+ MΦ
%F4/80+ (Out of total Live cells)
Control
Bifido MD
Desulfo
b
mLN HEV
Foxp3+ Treg
mLN HEV
F4/80+ MΦ
mLN HEV
CD11c+ DC
mLN CR
Foxp3+ Treg
mLN CR
F4/80+ MΦ
mLN CR
CD11c+ DC
MFI x % Area

### Suppl. Fig. S7

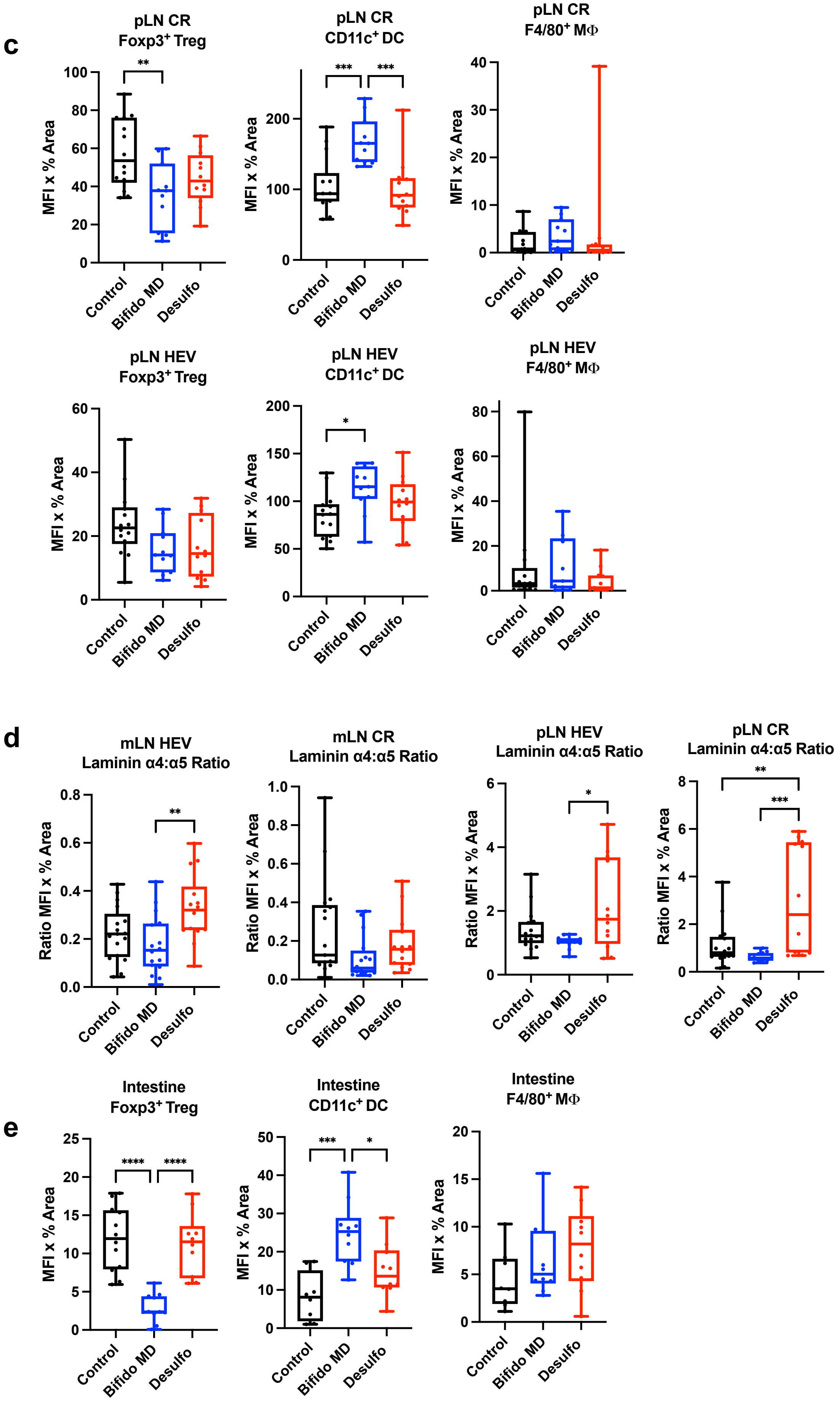

c
pLN CR Foxp3+ Treg
pLN CR CD11c+ DC
pLN CR F4/80+ MΦ
pLN HEV Foxp3+ Treg
pLN HEV CD11c+ DC
pLN HEV F4/80+ MΦ
MFI x % Area
Control
Bifido MD
Desulfo
d
mLN HEV Laminin α4:α5 Ratio
mLN CR Laminin α4:α5 Ratio
pLN HEV Laminin α4:α5 Ratio
pLN CR Laminin α4:α5 Ratio
Ratio MFI x % Area
e
Intestine Foxp3+ Treg
Intestine CD11c+ DC
Intestine F4/80+ MΦ
